# Life history effects on neutral diversity levels of autosomes and sex chromosomes

**DOI:** 10.1101/206862

**Authors:** Guy Amster, Guy Sella

## Abstract

All else being equal, the ratio of genetic diversity levels on X and autosomes at selectively neutral sites should mirror the ratio of their numbers in the population and thus equal ¾. Because X chromosomes spend twice as many generations in females as in males, however, the ratio of diversity levels is also affected by sex differences in life history. The effects of life history on diversity levels, notably those of sex-specific age structures and reproductive variances, have been studied for decades, yet existing theory relies on many parameters that are difficult to measure and lacks generality in ways that limit their applicability. We derive general yet simple expressions for these effects and show that life history effects on X-to-autosome (X:A) ratios of diversity levels depend only on sex-ratios of mutation rates, generation times, and reproductive variances. These results reveal that changing the sex-ratio of generation times has opposite effects on X:A ratios of polymorphism and divergence. They also explain how sex-specific life histories modulate the response of X:A polymorphism ratios to changes in population size. More generally, they clarify that sex-specific life history—generation times in particular—should have a marked effect on X:A polymorphism ratios in many taxa and enable the investigation of these effects.

**Significance Statement:** Understanding the determinants of neutral diversity patterns on autosomes and sex chromosomes provides a bedrock for our interpretation of population genetic data. Sex-specific age-structure and variation in reproductive success have long been thought to affect neutral diversity, but theoretical descriptions of these effects were complicated and/or lacked in generality, stymying attempts to relate diversity patterns of species with their life history. We derive general yet simple expressions for these effects, which clarify how they impact neutral diversity and should enable studies of relative diversity levels on the autosomes and sex chromosomes in many taxa.

## Introduction

Elucidating the forces that shape neutral diversity patterns has been a major obsession of modern population genetics (1–3) and a main driver of inferences from population genetic data (e.g., (4–7)). Contrasting relative neutral diversity levels on the X and autosomes is particularly interesting in this regard, as this contrast provides a unique setting for testing our understanding of the effects of evolutionary forces. All else being equal, the ratio of genetic diversity levels on X and autosomes should mirror the ratio of their numbers in the population and thus equal ¾. However, autosomes spend an equal number of generations in diploid form in both sexes, whereas the X spends twice as many generations in diploid form in females than in haploid form in males. As a result, the X to autosome (X:A) polymorphism ratio can also be shaped by sex differences in life history and mutation processes, as well as by differences in the effects of demographic history and selection at linked sites. (We focus on the X throughout, but similar arguments apply to neutral diversity on sex chromosomes more generally.)

While the effects of sex-specific age-structures on neutral diversity have been modeled for decades (8), these effects have been underappreciated in empirical studies of X:A ratios. Notably, in many species, generation times differ substantially between sexes (9), with likely implications for the polymorphism levels of X compared to autosomes. Indeed, parental ages are among the strongest known modifiers of mutation rates, and of the degree of male mutation bias (*α*) in mammals (10–14). Additionally, among mammals, longer male than female generation times lead to higher ratios of divergence levels on X versus autosomes on corresponding phylogenetic branches, presumably because they influence the relative number of generations that occurred on the X and autosomes (15). By the same token, we would expect sex-specific generation times to affect polymorphism ratios: for a given absolute time (in years) to the most recent common ancestor (MRCA) of X and autosome linked alleles, sex differences in generation times will lead to different numbers of generations on the X and autosomal lineages. In the case of polymorphism, however, this effect cannot be considered in isolation, as the coalescence process (and thus the distributions of times to the MRCAs on the X and autosomes) will also be affected by sex-specific generation times, and perhaps by sex specific age structure more generally (16).

The effects of age structure on the effective population size were first described by Felsenstein (8) in a haploid model, and later extended for the X and autosomes by others (e.g., (17–21)). Felsenstein assumed a haploid population, in which *M*_*i*_ individuals survive to age *i*, and a proportion *p*_*i*_ of newborns descend from parents of age *i*. He relied on identity by decent considerations to show that

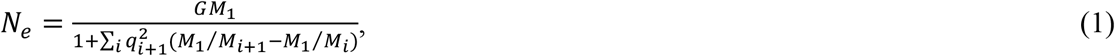

where *q*_*i*_ = Σ_*j*≥*i*_ *p*_*j*_, and *G* = Σ_*i*_ *i* · *p*_*i*_ is the expected generation time. While this result and its extensions to X and autosomes (17–21) provide valuable insights, they depend on a full parametrization of the age structure of the population. The dependence on so many parameters limits our understanding of the general effects of age structure on neutral diversity. Hill and Pollak derived alternative expressions for the *N*_*e*_ of autosomes (17) and of the X (21). For autosomes, for example, Hill showed that

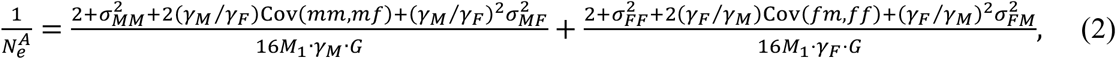

where 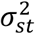 is the variance in the number of descendants of sex *t* that parents of sex *s* have throughout their life, Cov(*sm*, *sf*) is the covariance between the numbers of male (*sm*) and female (*sf*) descendants of parents of sex *s* throughout their life, and *γ*_*s*_ is the proportion of newborns of sex *s*. While Hill and Pollak’s expressions rely on fewer parameters, they are still quite complicated, again limiting our understanding of the effects of age structure. In both cases, the underlying parameters are also difficult to measure in nature, making it hard to relate the theoretical results with observed X:A polymorphism ratios. In contrast, the effects of a sex-specific age-structure on X:A ratios of neutral substitutions between species depend only on the sex ratio of generation times (15, 20), making them easy to understand and to test against phylogenetic observations (15).

Unlike sex-specific age structure, the effects of sex-specific variation in reproductive success on X:A polymorphism ratios have been considered by many empirical studies (e.g., (16, 22)). Notably, many species are known to be highly polygynous (i.e., a minority of males sire offspring with multiple females), implying higher reproductive variance in males than in females (16). Since the X spends twice as many generations in females than in males, higher male reproductive variance decreases coalescence rates on the X relative to on autosomes and thus increases the X:A polymorphism ratio. Both theoretical and empirical studies suggest that this effect can be substantial (e.g., (16, 22)).

From a theoretical standpoint, the effects of endogenous variation in reproductive success (endogenous as opposed to the variation introduced by stochastic birth and mortality) take simple forms when they are studied in isolation, i.e., without interactions with age structure. Notably, Wright (23) derived a simple expression for the effective population size in haploid models with endogenous reproductive variance and non-overlapping generations, i.e., without age-structure (see below); we present straightforward generalizations of this expression for X and autosomes (also see below). Since, several studies have incorporated reproductive variance into models with age structure, under the assumption that the age structure and reproductive variance are completely or partially independent (e.g., (24)). This assumption does not account for known correlations between ages of reproduction, reproductive success, and longevity: for example, between the age of first reproduction and longevity (25, 26) or between reproductive success and longevity (26, 27). A couple of studies considered more general models combining reproductive variance and age structure, but their analyses relied on simulations with particular parameter choices (28) or they resulted in complicated analytical results (16), with even more parameters than those with age-structure alone.

Our goal here is to understand how sex-specific age structure and variation in reproductive success affect neutral diversity levels on the X and autosomes in a general setting. Importantly, we aim to derive simple expressions for these dependencies, in terms of parameters that are straightforward to measure, so that the models can be related to and tested against observed diversity levels on the X and autosomes. To achieve these goals, we build on the coalescent treatment of age-structured populations (24, 29, 30).

## Results

### The haploid case

#### The Model

To illustrate how we treat the coalescence process in an age-structured population, we first consider the haploid model proposed by Felsenstein (8). We assume a haploid, panmictic population of constant size that is divided into age classes of one year (for convenience). We denote the number of individuals of age *a*_*max*_ ≥ *a* ≥ 1 by *M*_*a*_, where *M*_*a*_ is assumed to be constant, *M*_*a*+1_ ≤ *M*_*a*_ due to mortality, and *M*_*a*_ = 0 for ages *a* > *a*_*max*_ (see Table 1 for a summary of notations). We further assume that each of the *M*_1_ newborns is independently chosen to descend from a random parent: the ages of the parents are chosen from a distribution 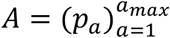 with expectation *G* (the generation time), and the specific parent is chosen with uniform probability within the age class (endogenous reproductive variance is considered below).

**Table 1.**
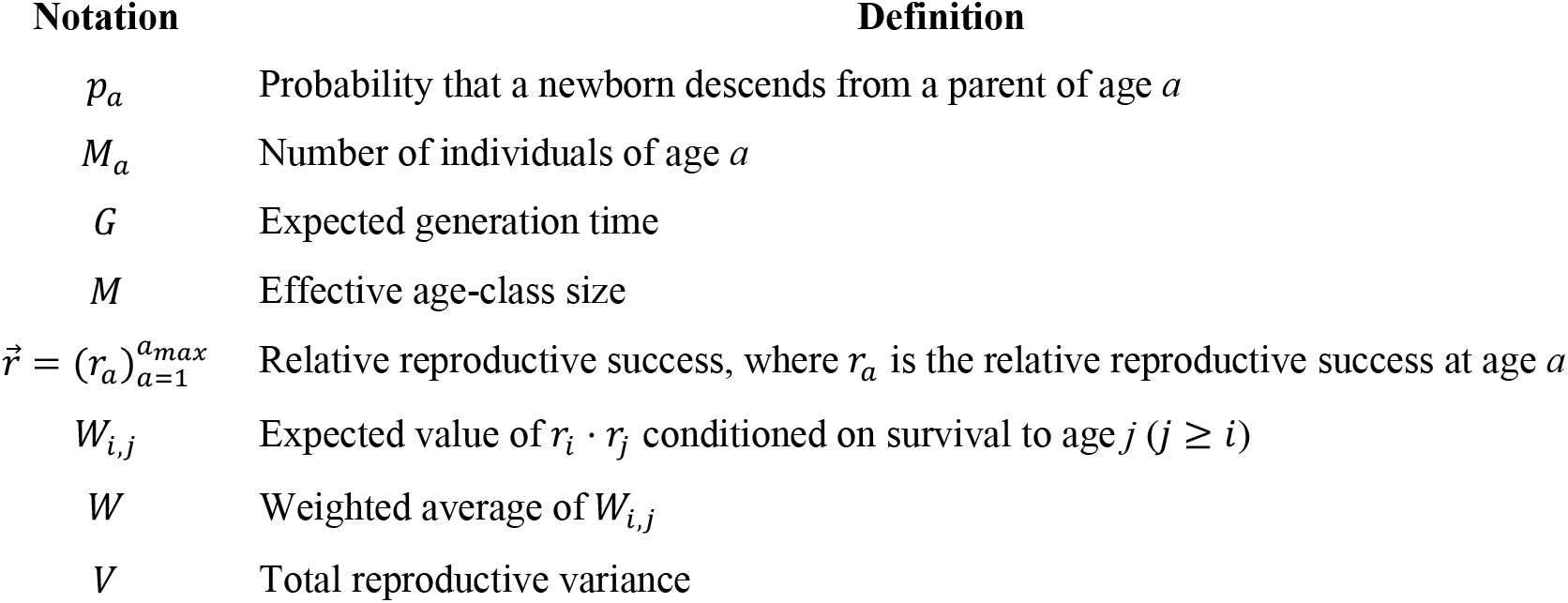
Notation for the haploid model.

| Notation | Definition |
| --- | --- |
| $p_a$ | Probability that a newborn descends from a parent of age $a$ |
| $M_a$ | Number of individuals of age $a$ |
| $G$ | Expected generation time |
| $M$ | Effective age-class size |
| $\vec{r} = (r_a)_{a=1}^{a_{max}}$ | Relative reproductive success, where $r_a$ is the relative reproductive success at age $a$ |
| $W_{i,j}$ | Expected value of $r_i \cdot r_j$ conditioned on survival to age $j$ ($j \geq i$) |
| $W$ | Weighted average of $W_{i,j}$ |
| $V$ | Total reproductive variance |

#### Effective population size

This model was solved by Sagitov and Jagers in a coalescent framework (24), and here we provide an intuitive account of the solution (see SI Section 1 for a rigorous derivation).

We first consider the rate of coalescence for a sample of two alleles; to this end, we trace their lineages backward in time. For the alleles to coalesce in a given age class *a*, the following conditions have to be met. One of them would have to be a newborn in the previous generation, which occurs with probability *p*_*a*_/*G* per year. The other allele would have to be present in the same age class, which occurs with probability Σ_*j*≥*a*_ *p*_*a*_/*G*, as it would have to be born to a parent of age *j* ≥ *a*, *j* − *a* + 1 generations ago (in SI Section 1.2 we show that this expression is in fact the stationary age distribution along a lineage). Finally, both alleles would also have to be in the same individual in age class *a*, which occurs with probability 1/*M*_*a*_. Thus, the probability of coalescence per year in age class *a* is

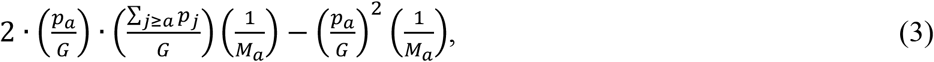

where we multiplied by 2, because either allele could have been the newborn in the previous generation, but subtracted the probability that they both were, because this event should be counted only once.

The probability of coalescence per generation and corresponding effective population size follow. Multiplying the probabilities per age class per year by the generation time, summing them over age classes, and rearranging terms, we find that

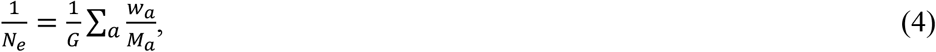

where 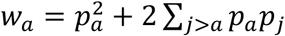. Here and throughout the paper, we follow standard coalescent theory in neglecting terms on the order of 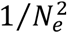 or smaller (31). Note that the (*a*, *j*)-term in *w*_*a*_ is proportional to the probability that the coalescence in age class *a* occurs in an individual that fathered a newborn carrying one of the alleles at age *a* and a newborn carrying the other allele at age *j*. The *w*_*a*_ terms add up to 1, allowing us to define the effective age class size, *M*, as the weighted harmonic mean

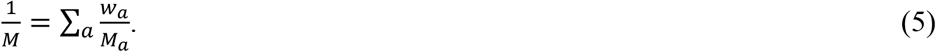

The effective population size can then be viewed as the product of the effective age-class size and the effective number of age-classes, which is simply the expected generation time *G*, i.e.,

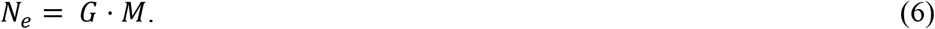

In SI Section 1.5, we show that Eq. 3 is equivalent to Felsenstein’s formula (Eq. 1 in (8)). The definition of the effective age-class size as a harmonic mean simplifies this formula and provides an intuition for the effect of age-structure on *N*_*e*_.

#### Reproductive variance

Next, we extend the model to incorporate endogenous reproductive variance. To this end, we assume that each newborn is assigned a vector 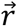 describing its age-dependent, relative reproductive success, such that its probability of being chosen as a parent among the individuals of age class *a* is *r*_*a*_/*M*_*a*_ (thus, *r*_*a*_ corresponds to the *expected*, rather than *realized*, reproductive success of the individual at age *a*). We further assume that the proportion of individuals with a given vector 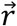 that reach age *a*, 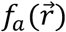, can vary with age, in effect allowing for dependencies between *expected* reproductive success and longevity. In SI Section 1.7, we detail why our results also apply to models that incorporate dependencies between *realized* reproductive success and longevity, such as the one proposed by Evans and Charlesworth (28). Our results thus apply to quite general interactions between reproductive success and age structure, including those that have been observed (see Introduction).

The extended model can be solved along the same lines we described above (see SI Section 1.2). Specifically, the coalescence rate per generation and corresponding effective population size take a similar form:

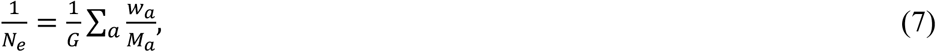

but in this case

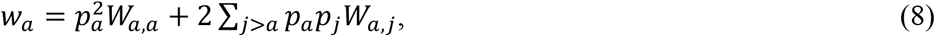

where 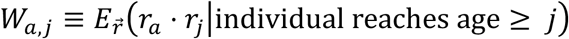 for *j* ≥ *a*. As in the simpler case, the (*a*, *j*)-term in *w*_*a*_ is proportional to the probability that coalescence in age class *a* occurs in an individual that fathered a newborn carrying one of the alleles at age *a* and a newborn carrying the other allele at age *j*; but in this case, the coefficients *w*_*a,j*_ incorporate the effect of endogenous reproductive variance. In contrast to the simpler case, the *w*_*a*_ terms do not necessarily add up to 1. We therefore introduce a normalization by *W* = Σ_*a*_ *w*_*a*_, and define the effective age class size as

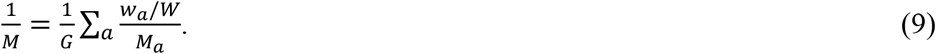

In these terms, the effective population size takes the form

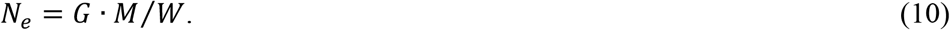

#### Recasting in terms of total reproductive variance

To provide further intuition, we first consider the special case in which relative reproductive success is independent of age and of mortality rates. Namely, each newborn is assigned a scalar relative reproductive success *r* at birth, such that its probability of being chosen as a parent among the individuals of age class *a* is *r*/*M*_*a*_; the distribution of *r* values then has expectation 1 (by construction). We denote its variance by *V*_*r*_(*V*_*r*_ = 0 when there is no endogenous reproductive variance). In this case, the coalescence rates in any given age class are increased by a factor 1 + *V*_*r*_, and therefore the effective population size is

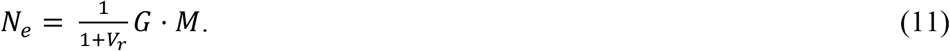

Thus, *W* = 1 + *V*_*r*_, which can be interpreted as the reproductive variance caused by the Poisson sampling of parents, which contributes 1, and by the endogenous reproductive variance, which contributes *V*_*r*_. In turn, the contribution of age-structure to reproductive variance is reflected in the term *G* · *M*.

The results of the general model can also be recast in terms of the total reproductive variance. First consider a haploid Wright-Fisher process (i.e., with non-overlapping generations) with endogenous reproductive variance, modeled similarly to the example considered above. In this case, Wright (23) showed that the effective population size is

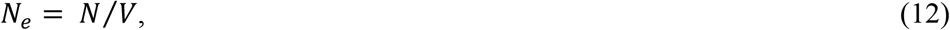

where *N* is the census population size and *V* = 1 + *V*_*r*_ is the total reproductive variance. Second, in the case with age-structure but without endogenous reproductive variance, Hill (17) showed that the effective population size can also be written as

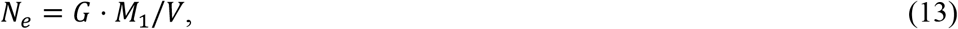

where *G* · *M*_1_ is the number of newborns per generation, and *V* is the reproductive variance introduced by age structure. Comparing Eqs. 6, 10 and 13, we see that this variance can be expressed as *V* = *M*_1_/*M* (≥ 1). This expression makes intuitive sense, as *V* defined in this way would be the reproductive variance in the case considered by Wright, if individuals in the next generation were randomly chosen from a reproductive pool including only *M* out of *M*_1_ individuals in the previous one. In SI Section 1.4, we show that Eq. 13 also holds for general models with age-structure and endogenous reproductive variance. In this case, the total reproductive variance is

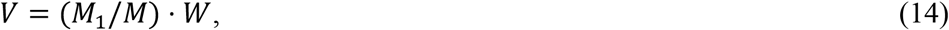

where the first term in this product, *M*_1_/*M*, is the variance introduced by stochasticity in birth and mortality, and the second term, *W*, reflects the contribution of endogenous reproductive variance.

In summary, Eq. 13 implies that all the effects of age-structure and endogenous reproductive variance (and any dependence between them) on the effective population size can be summarized in terms of the generation time, *G*, the number of newborns per year, *M*_1_, and the total reproductive variance, *V*. It also shows that, along with the total reproductive variance, it is the number of newborns per generation, *G* · *M*_1_, rather than the census population size, *N* = Σ_*a*_ *M*_*a*_, that determines the effective population size in age-structured populations. One implication is that (in the model with endogenous reproductive variance) *G* · *M*_1_ is an upper bound on *N*_*e*_ whereas *N* is not (see SI Section 1.6).

### X and autosomes

The diploid model with two sexes is more elaborate, but it is defined and solved along the same lines that we have described for the haploid model. Notably, in the diploid case, we allow for age-dependent mortality, fecundity, and endogenous reproductive variance to differ between the sexes. We also accommodate X and autosomal modes of inheritance (i.e., X linked loci in males always descend from females, whereas for autosomes the sex of parents is randomly chosen with probability ½). In SI Section 2.3, we solve this model for the stationary distributions of sex and age and corresponding coalescence rates, by extending Pollak’s results for age-structured populations with two sexes (30) to account for endogenous reproductive variance.

Here, we describe the two formulations for the effective population sizes on the X and autosomes, relying on the intuition gained from the haploid model. By analogy with the haploid case (Eq. 10), we can express the effective population sizes on the X and autosomes in terms of their respective effective sizes of age-classes. First, we define the basic quantities (*G*, *W*, and *M*) for each sex as in the haploid case. Second, we define these quantities for the X and autosomes as the appropriate averages over their values in males and females (Table 2). Formulated in this way, the effective population size for the X and autosomes take the form

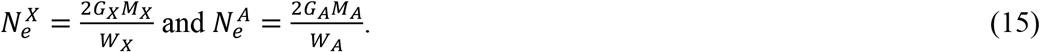

**Table 2.** Notation for the diploid model with two sexes. All quantities in males and females are defined as in the haploid model (Table 1). All quantities for X and autosomes, other than effective age-class sizes, are arithmetic averages over sexes, weighted by the proportion of generations that X and autosomes spend in each sex. The effective age-class sizes are harmonic averages over sexes, weighted by the relative endogenous reproductive success factors, *W*, in each sex.

| Notation | Definition |
| --- | --- |
| $M_1$ | Number of newborns (of both sexes) per-year |
| $\gamma_M, \gamma_F$ | Proportion of male and female newborns |
| $G_F, G_M$ | Expected generation times in females and males |
| $G_X = \frac{1}{3}G_M + \frac{2}{3}G_F,$<br>$G_A = \frac{1}{2}G_M + \frac{1}{2}G_F$ | Expected generation times for X and autosomes |
| $W_F, W_M$ | Expected endogenous reproductive success factors in males and females |
| $W_X = \frac{1}{3}W_M + \frac{2}{3}W_F,$<br>$W_A = \frac{1}{2}W_M + \frac{1}{2}W_F$ | Expected endogenous reproductive success factors for X and autosomes |
| $M_F, M_A$ | Effective sizes of age classes in males and females |
| $1/M_X = \frac{1}{3} \cdot (W_M/W_X)/M_M + \frac{2}{3} \cdot (W_F/W_X)/M_F,$<br>$1/M_A = \frac{1}{2} \cdot (W_M/W_A)/M_M + \frac{1}{2} \cdot (W_F/W_A)/M_F$ | Effective sizes of age classes for X and autosomes |
| $V_M, V_F$ | Total reproductive variance of males and females |
| $V_X = \frac{1}{3}V_M + \frac{2}{3}V_F,$<br>$V_A = \frac{1}{2}V_M + \frac{1}{2}V_F$ | Total reproductive variance for X and autosomes |
| $\mu_M, \mu_F$ | Expected mutation rate per generation in males and females |
| $\mu_X = \frac{1}{3}\mu_M + \frac{2}{3}\mu_F,$<br>$\mu_A = \frac{1}{2}\mu_M + \frac{1}{2}\mu_F$ | Average mutation rate on the X and autosomes |
| $f(x) = \frac{2x+4}{3x+3}$ | Function relating sex ratios and X:A ratios |

The factor 2, which is absent in the haploid case (Eq. 10), accounts for the effective number of age classes with two sexes (i.e., 2*G* instead of *G* classes in the haploid case). The corresponding coalescence rates per generation on X and autosomes are 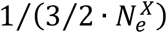 and 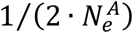, respectively. Note that sometimes the effective population sizes of X and autosomes are defined as the inverse of the coalescence rates (e.g., when saying that all else being equal, the X:A ratio of *N*_*e*_ is ¾) and sometimes they are defined as done here (e.g., such that they equal the number of individuals under the simple Wright-Fisher model).

We can also express the effective population sizes on the X and autosomes in terms of the number of newborns per generation and the total reproductive variances (by analogy with Eq. 13 for the haploid case). To this end, we first generalize Wright’s result for non-overlapping generations and endogenous reproductive variance (Eq. 12; (23)) to the X and autosomes. We assume that the population consists of *N*_*F*_ females and *N*_*M*_ males, with a total population size by *N* ≡ *N*_*M*_ + *N*_*F*_, and denote the proportion of each sex by *γ*_*s*_ ≡ *N*_*s*_/*N*, where *s* = *F*, *M*. Under this model

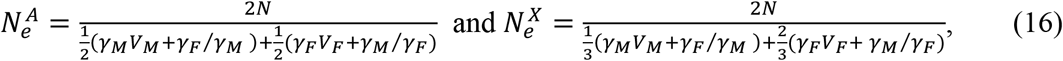

where *V*_*F*_ and *V*_*M*_ are the total reproductive variances in females and males, respectively (SI Section 2.4). With an equal sex ratio at birth (i.e., *γ*_*M*_ = *γ*_*F*_ = 1/2), these expressions reduce to

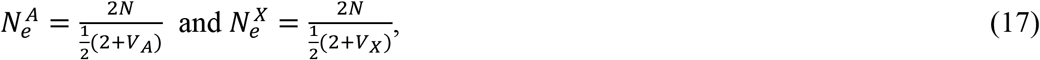

where *V*_*A*_ and *V*_*X*_ are the total reproductive variance on the X and autosomes, respectively (Table 2). Eqs. 16 and 17 differ from the analogous Eq. 12 for the haploid case in a couple of ways. In SI Section 2.6 we provide some intuition for these differences by recasting the effective population sizes on X and autosomes in terms of allelic rather than individual reproductive variances. Doing so makes the analogy between the haploid and diploid formulas apparent.

By analogy with the haploid case (Eq. 13), the expressions with general age-structure and endogenous reproductive variances follow from replacing the population size by the number of newborns per generation:

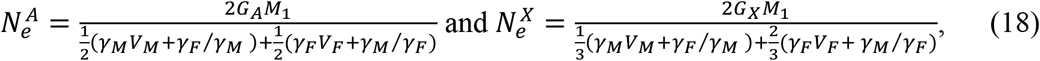

where in this case, the total variances *V*_*M*_ and *V*_*F*_ also include the effects of age-structure, *M*_1_ is the number of newborns of both sexes (per year), and the proportions of each sex, *γ*_*F*_ and *γ*_*M*_, are defined with respect to the numbers of newborns rather than individuals (SI Section 2.4). These expressions are considerably simpler than previous ones (see Introduction); in particular, they are given in terms of fewer parameters, which can be measured more readily in extant populations. Moreover, they demonstrate that the behavior under a general model of age-structure and endogenous reproductive variance is well approximated by the standard coalescence process (with non-overlapping generations and no endogenous reproductive variance) using the appropriate effective population sizes (Eq. 18) and units for time (i.e., using generation times on the X and autosomes, *G*_*X*_ and *G*_*A*_, respectively, to describe the coalescence process in years) (see SI Sections 1.2 and 1.3).

#### Polymorphism levels on X and autosomes

With expressions for the effective population sizes in hand, we turn to polymorphism levels. We allow for mutation rates to vary with sex and age (as they do in many taxa; (11)), with their per generation rates in females, *μ*_*F*_, and in males, *μ*_*M*_, defined as expectations over parental ages. The mutation rates per generation on the X, *μ*_*X*_, and autosome, *μ*_*A*_, are defined by the appropriate averages over their values in males and females (Table 2). The expected heterozygosity on X and autosomes then follow from the standard forms:

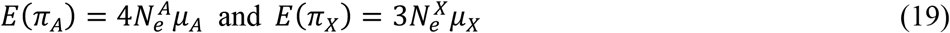

(1). Note that these expressions are usually derived assuming that the genealogical and mutational processes are independent (31). This assumption is violated here, because both the coalescence and mutation rates depend on the ages along a lineage. In SI Section 3, we show that the standard forms hold nonetheless.

We can now combine our results to derive expressions for X:A ratios. From Eq. 18, with some rearrangement of terms, the ratio of effective populations sizes is

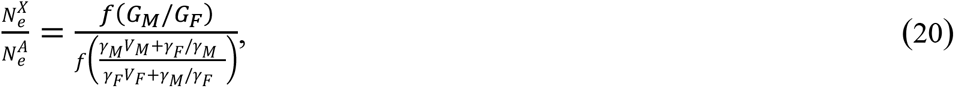

and from Eq. 19 the ratio of expected heterozygosities is

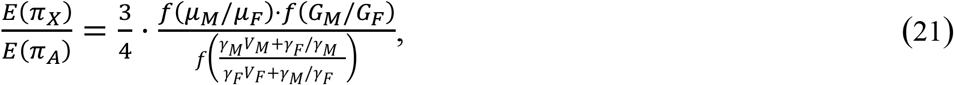

where 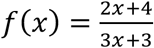 (given a quantity *C*, whose values on the X and autosomes are weighted arithmetic averages over sexes, 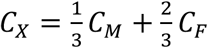 and 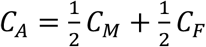 respectively, then *C*_*X*_/*C*_*A*_ = *f*(*C*_*M*_/*C*_*F*_)).

When mutation rates, age structures, and endogenous reproductive variances are identical in both sexes, Eq. 3 reduce to the naïve neutral expectation of ¾. When they vary between sexes, mutational and life history effects on the ratio reduce to the effects of the sex ratios of mutation rates (*α* = *μ*_*M*_/*μ*_*F*_), generation times (*G*_*M*_/*G*_*F*_), and reproductive variances (more precisely, (*γ*_*M*_*V*_*M*_ + *γ*_*F*_/*γ*_*M*_)/(*γ*_*F*_*V*_*F*_ + *γ*_*M*_/*γ*_*F*_)). Importantly, Eqs. 20 and 21 are much simpler and more general than previous results, and are expressed terms of sex ratios of parameters that are considerably easier to measure in extant populations.

## Discussion

Having a general yet simple expression for mutational and life history effects on the X:A polymorphism ratio allows us to draw several implications. First, it allows us to generalize previous bounds on the effects of each. Notably, we see that the multiplicative effect of the sex ratio of mutation rates and generation times is bound between 2/3 and 4/3 (see (32) for the mutational bound alone), and the effect of the sex ratio of reproductive variances is bound between 3/4 and 3/2 (see (16)). Considering these factors jointly, X:A polymorphism ratios are bound between ¼ and 2, a wider range than appreciated previously (note that these bounds apply only under a constant population size; see below). The functional dependence on the three sex ratios also suggests that changes in them will have the greatest impact when they are small (because |*f*′(*x*)| declines with *x*). As these sex ratios are male biased (i.e., greater than 1) in many taxa (33–35), we would expect differences in them among populations or closely related species to result in larger differences in X:A ratios when they are around 1, and to have much smaller effects if far from 1.

Second, our results clarify the differential effects of life history on X:A ratios of polymorphism and divergence. More precisely, we consider not divergence but the number of substitutions that accumulate on the X and autosomes on a lineage since a species split (i.e., ignoring multiple hits and the contribution of ancestral polymorphism), denoted *K*_*X*_ and *K*_*A*_ respectively. The equivalent expression to Eq. 2 for the ratio of substitutions is

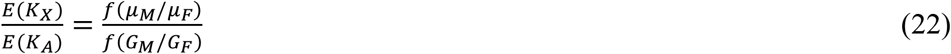

(15, 20). Thus, male mutation bias, *α* = *μ*_*M*_/*μ*_*F*_, has the same effect on ratios of polymorphism and substitutions. In contrast, reproductive variances and the sex ratio at birth affect only the polymorphism ratio because they affect the relative rates of coalescence and thus the relative lengths of lineages on the X and autosomes; for the substitutions ratio, this length is set by the species’ split time. Interestingly, the sex ratio of generation times (*G*_*M*_/*G*_*F*_) has opposite effects on polymorphism and substitutions ratios (Fig. 1). These opposing effects arise because generation times affect not only the relative number of generations on X compared to autosomes, which affects both polymorphism and substitution ratios, but also relative coalescence rates, which affects only the polymorphism ratio.

**Figure 1.**
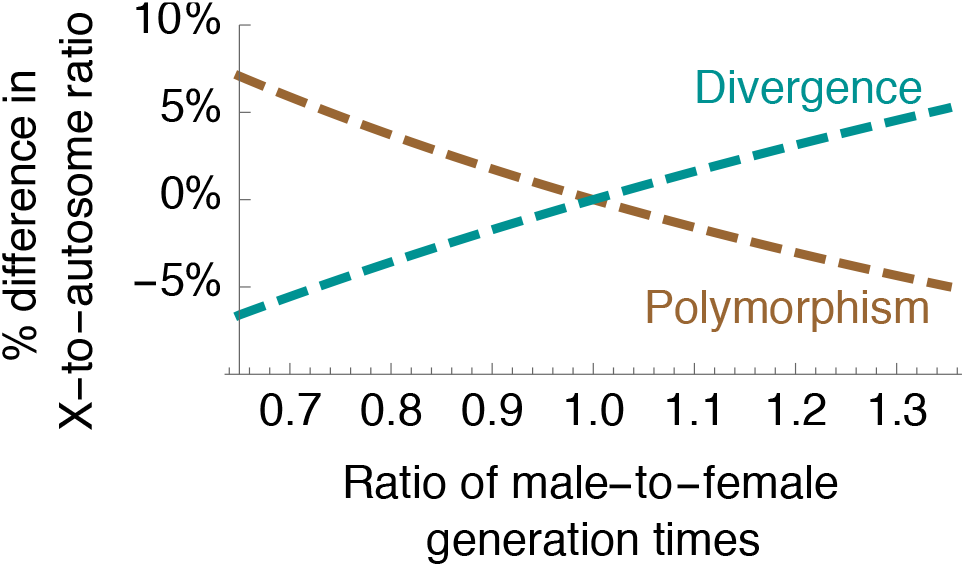
The sex ratio of generation times (*G*_*M*_/*G*_*F*_) has opposite effects on X:A ratios of polymorphism levels and substitutions.

Third, we can rely on the results for a constant population size to study how changes in population size and life history over time jointly affect X:A polymorphism ratios. Changes in population size are known to affect patterns of genetic variation in general (this signal underpins modern demographic inference, e.g., (4)), and X:A polymorphism ratios in particular (36–38). Having shown that life history also dramatically affects X:A ratios, it is natural to ask how these effects act jointly. In SI Section 4, we extend our model to incorporate these effects. We assume that population size and sex-specific life histories and mutation rates are piecewise-constant in *n* time intervals, where the *i*-th interval moving backwards from the present is [*T*_*i*−1_, *T*_*i*_), and 0 = *T*_0_ < *T*_1_ < ⋯ < *T*_*n*_ = ∞. We show that the expected heterozygosity on autosomes at the present is well approximated by the recursion

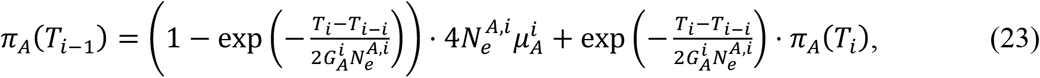

where 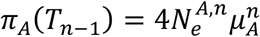, and heterozygosity on the X follows a similar recursion (Eqs. S136 and S137). The ratio 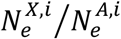 in any given interval *i* follows from the effects of the sex ratios of generation times and reproductive variances in a constant population size (Eq. 20). These recursions can be solved for *π*_*A*_ and *π*_*X*_ at the present and for their values at any time in the past (i.e., by substituting *t* for *T*_0_ in the recursions). In SI Section 4.3, we further describe how standard coalescence simulators can be used to incorporate the effects of sex- and time-dependent mutation rates, reproductive variances and age structure.

To illustrate how life history modulates the effects of changes in population size on the X:A ratio, we consider a simple scenario, in which the autosomal population size drops from *N*_*A*_ to *k* · *N*_*A*_ at time *t* = 0, where here we consider *t* to be increasing forward in time. We further assume that sex ratios of generation times, reproductive variances, and mutation rates are constant, and set the ratio of mutation rates to *α* = 1 to isolate genealogical effects. Under these assumptions, the equilibrium X:A polymorphism ratio, which is also the ratio at times *t* < 0, is 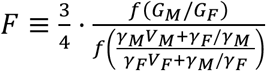 (with *α* ≠ 1, *F* is further multiplied by *f*(*α*)). Solving the recursions (Eqs. 23 and S137), we find that the polymorphism ratio at times *t* ≥ 0 is

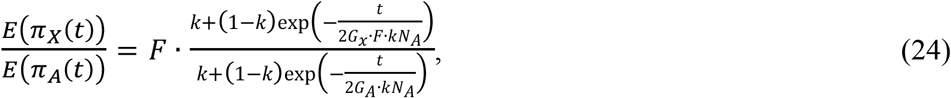

which converges to the equilibrium ratio *F* in the long run (i.e., when *t* ≫ *G*_*A*_ · *k* · *N*_*A*_).

This example illustrates how life history modulates the transient response of the X:A ratio to changes in population size (Fig. 2). Without sex-specific life histories, i.e., when *F* = 3/4 and *G* ≡ *G*_*X*_ = *G*_*A*_, Eq. 4 reduces to the expression derived by Pool and Nielsen (37). As they show, higher coalescence rate on the X compared to autosomes result in faster equilibration of X-linked diversity levels to the new population size, with exponential rate [3/4 · (2*G* · *kN*_*A*_)]^−1^ compared to [2*G* · *kN*_*A*_]^−1^, and thus in a transient reduction of the X:A polymorphism ratio (black curve in Fig. 2). A higher male reproductive variance weakens the difference between X and autosome (blue curve in Fig. 2), because it causes coalescence rates (and thus rates of equilibration) to be more similar on the X and autosomes, decreasing equilibration rates on the X by a factors of 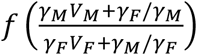. On the other hand, a longer male generation time enhances the difference between X and autosomes in two ways (red curve in Fig.2): it increases coalescence rates and thus equilibration rates *per generation* on the X compared to autosomes (by a factor of 1/*f*(*G*_*M*_/*G*_*F*_)), and it also decreases the relative *number of generations* that elapsed since the drop in population size on the X compared autosomes (by another factor of [*G*_*X*_/*G*_*A*_]^−1^ = 1/*f*(*G*_*M*_/*G*_*F*_)). More generally, sex-specific life history modulates the effects of changes in population size on X:A polymorphism ratios in two ways: one is by changing the relative X:A coalescence time scale of the response *in generations*; the other is by changing the relative X:A generation times, and thus the relative rate of response of X versus autosome *per unit time* (in years).

**Figure 2.**
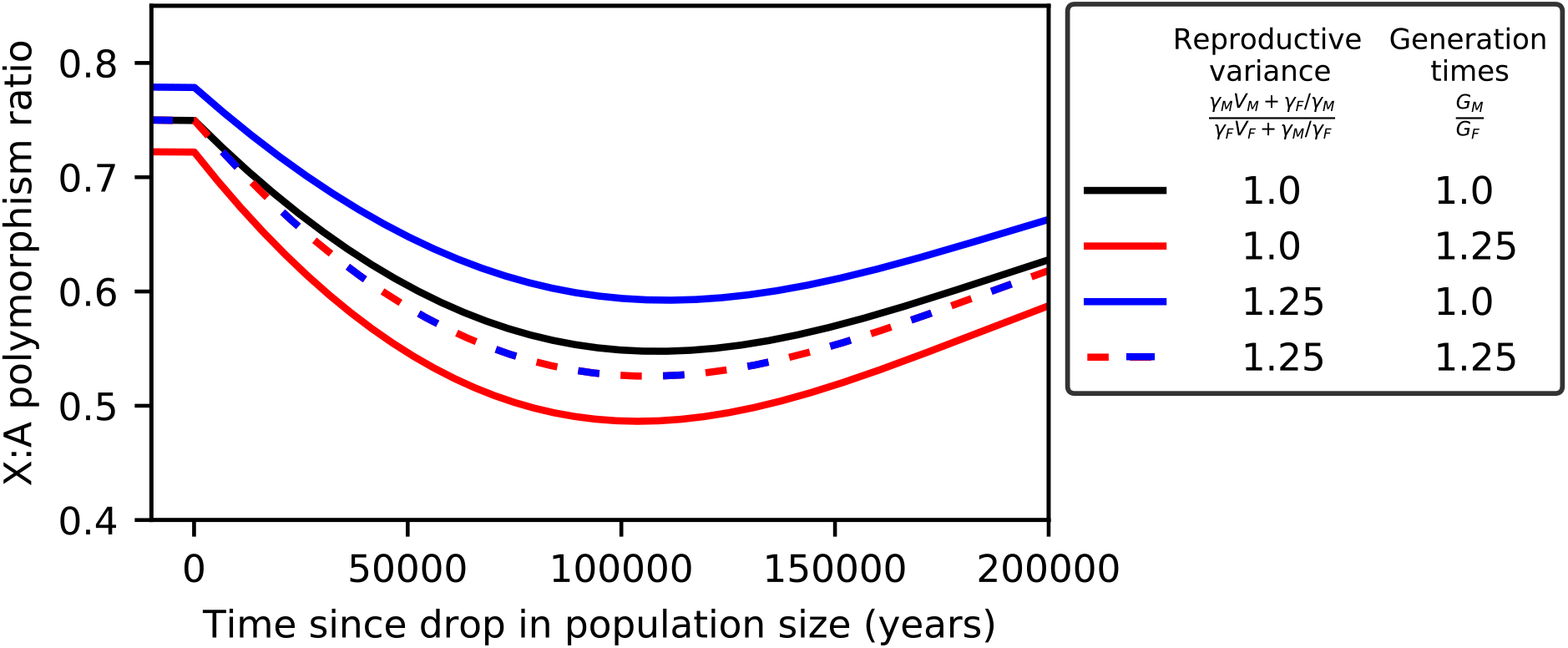
Sex-specific life histories modulate the responses to a change in population size on the X and autosomes. We consider a scenario in which the autosomal population size dropped from *N*_*A*_ = 10,000 to *N*_*A*_ = 1000 at time *t* = 0, and show the response of the X:A polymorphism ratio afterwards. We assume an autosomal generation time of 30 years, a sex-ratio of mutation rates of *α* = 1, with the reproductive variances and generation times ratios specified in the Figure.

Fourth, our results indicate that age-structure likely has underappreciated and non-negligible effects on the genetic diversity of X (or Z) and autosomes in many species. Notably, sex-specific generation times are the strongest known modifier of mutation rates (10, 11, 39) and, as we show, their genealogical effects on divergence and polymorphism on autosomes and sex chromosomes can also be substantial. Our results can therefore help to interpret observed polymorphism ratios in many species, given estimates for life history trait values. In particular, in a parallel paper (40), we take such an approach to show that after accounting for the differential effects of linked selection on the X and autosomes, the joint effects of changes in life history and population size can fully explain the variation in X:A polymorphism ratios across human populations.

## Acknowledgements

We thank M. Nordborg for helpful discussions and M. Przeworski for many helpful discussions and comments on the manuscript. We also thank the editor and three anonymous reviewers for many helpful comments on an earlier version of this manuscript.

