## Supplemental Information for "Life history effects on neutral diversity levels of autosomes and sex chromosomes"

### 1. Haploid Model

Here we rigorously solve the haploid model with age-structure and endogenous reproductive variance, relate our results to previous work that considered special cases, investigate the properties of the effective population size in age-structured populations, and show that our main results also apply for general dependencies between realized reproduction at different ages. In Section 1.1 we spell out our assumptions about endogenous reproductive variance. In Section 1.2 we solve for the joint stationary distribution of the age and relative reproductive success associated with an allele, going backwards in time. Based on this distribution, we calculate the stationary per-generation coalescence rate for a sample of two alleles, to obtain Eq. 10 in the main text:

$$N_e = \frac{M \cdot G}{W}. \tag{S1}$$

In Section 1.3 we formally show that Eq. 10 holds in the limit in which the census population size goes to infinity while the population's age structure is held constant; we then derive a general and tight bound on the rate at which this solution is approached as the population size is increased. In Section 1.4, we recast our results in terms of total reproductive variance, to show that the relationship derived by Hill for the case with age-structure alone (1):

$$N_e = G \cdot M_1/V, \tag{S2}$$

which is our Eq. 13, applies to the extended model with endogenous reproductive variance. We also show that the total reproductive variance in this case is

$$V = W \cdot (M_1/M), \tag{S3}$$

which is our Eq. 13. This concludes the derivations of our main results.

In Section 1.5 we show that in the case without endogenous reproductive variance, our Eq. 10 (Eq. S4 above) reduces to Felsenstein's formula (2), and consider a simple example of how age-structure affects the effective population size. In Section 1.6 we investigate the properties of the effective population size in age-structured populations with endogenous reproductive variance. In particular, we derive an upper bound for the effective population size and derive the conditions under which it can be attained. In Section 1.7, we consider an alternative model, which allows for general dependencies between realized (rather than expected) reproductive success at different ages, and show that Hill's formula also applies to this model.

#### 1.1 Requirements on $f_a$

When we introduced the haploid model with endogenous reproductive variance, we assumed that each newborn is assigned a relative reproductive success vector $\vec{r}$, where the (constant) proportion of individuals with a given vector $\vec{r}$ in age class $a$ was denoted by $f_a(\vec{r})$ (see Table S1 for summary of notation). Here we describe the requirements on the probability mass function $f_a$ that these assumptions entail. First, given that the probability of being born to a parent of age $a$ is $p_a$, and to a specific parent of age $a$ and with reproductive success $\vec{r}$ is $p_a \cdot \frac{r_a}{M_a}$, we require that $E_{f_a}(r_a) = \sum_{\vec{r}} f_a(\vec{r}) \cdot r_a = 1$ for any age $a$. Second, given that the number of individuals with a given $\vec{r}$ can only decrease with age (due to mortality), we further require that $M_a \cdot f_a(\vec{r}) \geq M_{a+1} \cdot f_{a+1}(\vec{r})$.

Third, requiring that the number of individuals of a given age $a$ and with a given $\vec{r}$ is constant and equal to $M_a \cdot f_a(\vec{r})$ implies that this number needs to be an integer. Notably, if we would like to model the distribution of relative reproductive success using a given (continuous or discrete) distributions $\tilde{f}_a(\vec{r})$, which satisfies the first two requirements, we would need to discretize $\tilde{f}_a$ to obtain a probability mass function $\tilde{f}'_a$ such that $M_a \cdot \tilde{f}'_a(\vec{r})$ is always an integer. However, if we assume that the relative sizes of the age-class, i.e., the ratios $M_i/M_j$, are constant, and increase the total population sizes, the discretized functions $\tilde{f}'_a$ will approach $\tilde{f}_a$, and the value of the $W_{i,j} = E_{\tilde{f}'_j}(r_i \cdot r_j) = E(r_i \cdot r_j | \text{survival to age } j)$ terms, which summarize the effect of endogenous reproductive variance on the effective population size, will approach $E_{\tilde{f}_j}(r_i \cdot r_j)$. We implicitly assumed this limit when we considered the special case in which relative reproductive success is independent of age and of mortality rates. More generally, while the assumption that for any age $a$, $M_a \cdot f_a(\vec{r})$ is an integer, might appear highly restrictive, these restrictions are relaxed under the standard coalescent assumption that the population size is sufficiently large.

| Notation | Definition | Remarks |
|---|---|---|
| $p_a$ | Probability that a newborn descends from a parent of age $a$ | $\sum_a p_a = 1$ |
| $q_a$ | Probability that a newborn descends from a parent of age $\geq a$ | $q_a = \sum_{i \geq a} p_i$ |
| $G$ | Expected generation time | $G = \sum_a a \cdot p_a$ |
| $M_a$ | Number of individuals of age $a$ ($M_1$ is the number of newborns per-year) | $M_{a+1} \leq M_a$ |
| $\vec{r}$ | Relative reproductive success, where component $r_a$ is the relative reproductive success at age $a$ | |
| $f_a(\vec{r})$ | The proportion of individuals with relative reproductive success $\vec{r}$ among individuals of age $a$ | |
| $g_a(\vec{r})$ | Given an individual $I$ of age $a$ and a newborn $n$, $g_a(\vec{r})$ is the probability that $I$ has relative reproductive success $\vec{r}$, conditioned on $n$ being descended from $I$ | $g_a(\vec{r}) = r_a f_a(\vec{r})$ |
| $\epsilon(a, \vec{r})$ | Joint stationary probability of age $a$ and relative reproductive success $\vec{r}$ along a lineage, going backwards in time | $\epsilon(a, \vec{r}) = \frac{1}{G} \sum_{j \geq a} p_j g_j(\vec{r})$ |
| $\epsilon_a$ | Marginal stationary distribution of age $a$ | $\epsilon_a = \frac{q_a}{G}$ |
| $M$ | Effective age-class size | See Eq. S17 |
| $W_{i,j}$ | Average value of $r_i \cdot r_j$ among individuals of sex $s$ and age $a$ | Defined for $i \leq j$ |
| $W$ | Weighted average of the $W_{i,j}$ | See Eq. S16 |
| $X, X_a$ | An individual's number of offspring, throughout its life or at age a, respectively | |
| $V$ | Reproductive variance (i.e., $V = Var(X)$) | See Eq. S44 |
| $S_a$ | The event of surviving to age $\geq a$ | |

**Table S1:** Notation for the haploid model, with parameters of the model in red.

#### 1.2 Stationary coalescence rate and effective population size

Here, we extend the derivations of Sagitov and Jagers (3) to account for endogenous reproductive variance. Tracing an allele backward in time, the age $a_t$ and relative reproductive success $\vec{r}_t$ of the individual $I_t$ who carries the allele $t$ years in the past defines a Markov chain $(a_t, \vec{r}_t)$. To define the transition probabilities of the chain, we distinguish between two cases. First, if the individual carrying the allele is not a newborn, i.e., $a_t > 1$, then at time $t$+1 that individual will be one year younger and its relative reproductive success $\vec{r}$ will remain unchanged, i.e., $(a_{t+1}, \vec{r}_{t+1}) = (a_t - 1, \vec{r}_t)$ with probability one. Second, if the individual carrying the allele is a newborn, i.e., $a_t = 1$, then $a_{t+1}$ equals $a$ with probability $p_a$. The probability mass function of $\vec{r}_{t+1}$ conditional on $a_{t+1}$, follows from Bayes' theorem, further conditioning on the fact that the parent, $I_{t+1} = I$, necessarily reproduced successfully

$$P(\vec{r}_I = \vec{r} | I_{t+1} = I, a_{t+1} = a)$$
$$= \frac{P(I_{t+1}=I|\vec{r}_{t+1}=\vec{r}, a_{t+1}=a) \cdot P(\vec{r}_I = \vec{r}|\, a_{t+1}=a)}{P(I_{t+1}=I)} = \frac{(r_a/M_a) \cdot f_a(\vec{r})}{\sum_{\vec{k}} (r_a/M_a) \cdot f_a(\vec{k})} = r_a \cdot f_a(\vec{r}). \quad \text{(S5)}$$

We denote this probability by $g_a(\vec{r}) \equiv r_a \cdot f_a(\vec{r})$, and conclude that

$$P((a_{t+1}, \vec{r}_{t+1}) = (a, \vec{r}) | a_t = 1) = p_a \cdot g_a(\vec{r}). \quad \text{(S6)}$$

$g_a$ is a proper probability mass function since $\sum_{\vec{r}} g_a(\vec{r}) = \sum_{\vec{r}} r_a \cdot f_a(\vec{r}) = 1$. Moreover, the parent's expected value of $r_a$ is $E_{\vec{r} \sim g_a}(r_a) = E_{\vec{r} \sim f_a}(r_a^2) = 1 + V_{\vec{r} \sim f_a}(r_a) \geq 1$. The latter inequality makes intuitive sense, as it implies that the allele is more likely to be descended from an individual that has higher than average relative reproductive success in its age class.

We rely on the transition probabilities to derive and solve a recursion for the stationary probability $\epsilon(a, \vec{r})$ of age, $a$ and relative reproductive successes, $\vec{r}$, of the individuals carrying the allele. Namely,

$$\epsilon(a, \vec{r}) = \epsilon(a + 1, \vec{r}) + \left(\sum_{\vec{k}} \epsilon(1, \vec{k})\right) \cdot p_a \cdot g_a(\vec{r}), \quad \text{(S7)}$$

where the first term corresponds to aging within the same individual and the second corresponds to parenting a newborn. In order to solve these recursions, we first consider the marginal stationary distribution of age, $\epsilon_a = \sum_{\vec{r}} \epsilon(a, \vec{r})$. To this end, we sum the recursions over $\vec{r}$ to obtain recursions on the marginal distribution,

$$\epsilon_a = \epsilon_{a+1} + \epsilon_1 \cdot p_a, \quad \text{(S8)}$$

where we also require that $\sum_a \epsilon_a = 1$. This recursion was solved by Sagitov and Jagers (3) for the case without endogenous reproductive variance, yielding

$$\epsilon_a = q_a / G, \quad \text{(S9)}$$

where $q_a \equiv \sum_{j \geq a} p_j$. Substituting this expression into Eq. S7, the recursions simplify to

$$\epsilon(a, \vec{r}) = \epsilon(a + 1, \vec{r}) + \frac{1}{G} \cdot p_a \cdot g_a(\vec{r}), \quad \text{(S10)}$$

where we further require that $\sum_{a,\vec{r}} \epsilon(a,\vec{r}) = 1$. The solution of these recursions is

$$\epsilon(a,\vec{r}) = \frac{1}{G}\sum_{j\geq a} p_j g_j(\vec{r}). \quad \text{(S11)}$$

The marginal stationary probability mass function of $\vec{r}$ is $\sum_a \epsilon(a,\vec{r}) = \frac{1}{G}\sum_j (j \cdot p_j) \cdot g_j(\vec{r})$, which is a proper probability mass function because $\frac{1}{G}\sum_j j \cdot p_j = 1$, and $\sum_{\vec{r}} g_a(\vec{r}) = 1$ for any age $a$.

We rely on the stationary distribution to derive the probability of coalescence of two alleles, along the same lines as detailed in the main text for the case without endogenous reproductive variance. For the coalescence to occur at time *t* in the past, one of the alleles (A) would descend from the other (B) or both would descend from the same parental allele at that time (this is contrary to the case of non-overlapping generations, in which coalescence necessarily occurs when both alleles descend from the same parental allele in the previous generation). For example, if allele A is associated with a newborn, one coalescence scenario would be for allele B to be associated with an individual of age $a > 1$, from which A descends a single time step (e.g., year) further in the past; a second coalescence scenario would be for both alleles to be associated with newborns at the same time, and descend from the same individual. Specifically, if allele B is in an individual of age *a* and relative reproductive success $\vec{r}$ at time *t* (with probability $\epsilon(a,\vec{r})$), then allele A must be in a newborn at time *t*-1 (with probability $\epsilon_1$) having descended from the same individual carrying allele B (with probability $p_a \cdot \frac{r_a}{M_a}$). Summing over the individual's possible ages and reproductive success vectors, we obtain the probability

$$\sum_{a,\vec{r}} \epsilon(a,\vec{r}) \cdot \epsilon_1 \cdot p_a \cdot \frac{r_a}{M_a} = \frac{1}{G^2}\sum_a \frac{\sum_{j\geq a} p_a p_j \sum_{\vec{r}} r_a g_j(\vec{r})}{M_a} = \frac{1}{G^2}\sum_a \frac{\sum_{j\geq a} p_a p_j W_{a,j}}{M_a}, \quad \text{(S12)}$$

where for $j \geq i$,

$$W_{i,j} = \sum_{\vec{r}} r_i g_j(\vec{r}) = \sum_{\vec{r}} r_i r_j f_j(\vec{r}) = E_{\vec{r}\sim f_j}(r_i \cdot r_j) \quad \text{(S13)}$$

is the expectation of $(r_i \cdot r_j)$ conditional on surviving to age $\geq j$. Further allowing for either allele or both to be the newborn (using the inclusion-exclusion principal to subtract the probability $\epsilon_1^2 \sum_a \frac{p_a^2 W_{a,a}}{M_a}$ that both alleles were in a newborn prior to coalescence), and measuring the coalescence rate in generations (rather than years), we obtain the per-generation coalescence rate and corresponding effective population size:

$$\frac{1}{N_e} = \frac{1}{G}\sum_a \frac{p_a^2 W_{a,a} + 2\sum_{j>a} p_a p_j W_{a,j}}{M_a}. \quad \text{(S14)}$$

Eq. S14 can be rearranged to obtain Eq. 7 of the main text. To this end, we define

$$w_i = (p_i^2 W_{i,i} + 2 \sum_{j>i} p_i p_j W_{i,j})/W, \tag{S15}$$

where

$$W = \sum_i p_i^2 W_{i,i} + 2 \sum_{i<j} p_i p_j W_{i,j} \tag{S16}$$

is a weighted average of the $W_{i,j}$. Noting that $\sum_a w_a = 1$, we then define the effective age class size as a weighted harmonic average of the age class sizes,

$$\frac{1}{M} = \sum_a \frac{w_a}{M_a}. \tag{S17}$$

Substituting this expression into Eq. S14, we obtain Eq. 7 of the main text:

$$\frac{1}{N_e} = W/(M \cdot G). \tag{S18}$$

#### 1.3 Convergence to the stationary solution

Here, we provide a formal justification for using the stationary coalescence rate and effective population size. To this end, we define the coalescence process and expected *TMRCA* rigorously, detail the condition under which our asymptotic expression for the expected *TMRCA* and thus effective population size (Eq. 10 in the main text) are exact, and derive a tight bound on the rate of convergence to the asymptotic rate of coalescence as the population size increases. Lastly, we relate our results to previous work that considered the convergence to asymptotic coalescence rates.

In Section 1.2 we modelled the state of an allele as a Markov chain. We defined the state space of the chain as the set $S$ containing all possible pairs $s = (a, \vec{r})$, where $a$ is the age of the allele, and $\vec{r}$ its relative reproductive success. We calculated the transition matrix, in which $P(s,t)$ is the transition probability between state $s$ and state $t$, and its stationary distribution $\epsilon(s) = \epsilon(a, \vec{r})$. In doing so and in what follows, we assume the existence of a unique stationary distribution $\epsilon$. While this requirement imposes conditions on model parameters, we expect it to be satisfied for realistic age-structures. Notably, the non-trivial requirement is for the chain to be aperiodic, which would be satisfied, for example, if there are two consecutive ages in which reproduction can occur (4).

To define the coalesce process of two alleles formally, we first need to define the alleles' joint states as a Markov chain. We do so in two steps. First, we consider a chain $X_t^\infty$ in which the states of the alleles are completely independent. This definition corresponds to a hypothetical infinite population with the specified age-structure, in which the alleles never coalesce. The state space of the chain is $S^{\infty} = S \times S$, and its transition matrix is $P^{\infty}(\vec{s}, \vec{t}) = P(s_1, t_1) \cdot P(s_2, t_2)$, with subscripts corresponding to each of the alleles. It follows that the stationary distribution associated with this chain is $\pi^{\infty}(s_1, s_2) = \epsilon(s_1) \cdot \epsilon(s_2)$.

Second, we consider a corresponding chain $X_t^M$ that incorporates coalescence. This chain corresponds to the same age-structure, but with a finite effective age class size $M$. The state space of the chain is $S^M = S^{\infty} \cup \{c\}$, where $c$ is an absorbing state, to which the chain transitions upon coalescence (we do not track the specific state of the allele after coalescence). The transition matrix of this chain for $\vec{s}, \vec{t} \in S^{\infty}$ is:

$$P^M(\vec{s}, \vec{t}) = P^{\infty}(\vec{s}, \vec{t}) \cdot (1 - C_{\vec{s},\vec{t}}),$$

$$P^M(c, \vec{t}) = 0,$$

$$P^M(c, c) = 1, \text{ and}$$

$$P^M(\vec{s}, c) = \sum_{\vec{y} \in S \times S} P^{\infty}(\vec{s}, \vec{y}) \cdot C_{\vec{s},\vec{y}}, \qquad \text{(S19)}$$

where $C_{\vec{s},\vec{t}}$ is the probability of coalescence conditioned that the two alleles transition from states $\vec{s}$ to states $\vec{t}$. Note that the stationary distribution of the chain $X_t^M$ is trivial, i.e., it is $c$ with probability 1 (and not $\pi^{\infty}$).

The probabilities of coalescence $C_{\vec{s},\vec{t}}$ can be described explicitly using the parameters of the model, but we do not require the explicit form for our purposes here. What is of interest to us, as will become clear below, is their weighted average:

$$\sum_{\vec{x},\vec{y} \in S^{\infty}} \pi^{\infty}(\vec{x}) \cdot P^{\infty}(\vec{x}, \vec{y}) \cdot C_{\vec{x},\vec{y}}. \qquad \text{(S20)}$$

This average is the probability that a pair of alleles, each drawn independently from the stationary distribution $\epsilon$, coalesce in some state y within a single time step. In Section 1.2 we referred to this probability informally as the 'stationary probability of coalescence per-year', and found it to be

$$\sum_{\vec{x},\vec{y} \in S^{\infty}} \pi^{\infty}(\vec{x}) \cdot P^{\infty}(\vec{x}, \vec{y}) \cdot C_{\vec{x},\vec{y}} = \frac{W}{M \cdot G^2}. \qquad \text{(S21)}$$

Next, we define the *TMRCA* of a sample of two alleles. Using the standard definition of a hitting time, i.e., $\tau_A = \min\{t \geq 0 : X_t \in A\}$, the *TMRCA* of the sample is simply $\tau_{\{c\}}^M$, where by $\tau^M$ and

$\tau^{\infty}$ we refer to the hitting times of the chains $X_t^M$ and $X_t^{\infty}$, respectively. We denote the expected *TMRCA*, given an initial state of the chain $X_0^M \in S^{\infty}$, by

$$e_x \equiv E(\tau_{\{c\}}^M | X_0^M = x). \quad \text{(S22)}$$

Note that this definition is restricted to the case in which the two alleles are initially distinct (i.e., $X_0^M \neq c$).

We can now derive several results about the expected *TMRCA*. By conditioning on the first step, i.e., on the value of $X_1^M = y$, we find that

$$e_x = 1 + \sum_{y \in S^{\infty}} P^M(x,y) \cdot e_y = 1 + \sum_{y \in S^{\infty}} P^{\infty}(x,y) \cdot (1 - C_{x,y}) \cdot e_y. \quad \text{(S23)}$$

It follows that if the initial state of the chain is sampled from the stationary distribution of the chain $X_t^{\infty}$ then the expected *TMRCA* satisfies

$$\sum_{x \in S^{\infty}} \pi^{\infty}(x) \cdot e_x = 1 + \boldsymbol{\sum_{x,y \in S^{\infty}} \pi^{\infty}(x) \cdot P^{\infty}(x,y) \cdot e_y} - \sum_{x,y \in S^{\infty}} \pi^{\infty}(x) \cdot P^{\infty}(x,y) \cdot C_{x,y} \cdot e_y. \quad \text{(S24)}$$

The term in bold, simplifies to

$$\sum_{x,y \in S^{\infty}} \pi^{\infty}(x) \cdot P^{\infty}(x,y) \cdot e_y = \sum_{y \in S^{\infty}} [\sum_{x \in S^{\infty}} \pi^{\infty}(x) \cdot P^{\infty}(x,y)] \cdot e_y = \sum_{y \in S^{\infty}} \pi^{\infty}(y) \cdot e_y, \quad \text{(S25)}$$

and therefore Eq. S24 simplifies to

$$\sum_{x,y \in S^{\infty}} \pi^{\infty}(x) \cdot P^{\infty}(x,y) \cdot C_{x,y} \cdot e_y = 1. \quad \text{(S26)}$$

Relying on Eq. S21, we can then rewrite Eq. S26 as

$$\sum_{y \in S^{\infty}} \alpha_y \cdot e_y = \frac{M \cdot G^2}{W}, \quad \text{(S27)}$$

where $\alpha(y) = \frac{\sum_{x \in S^{\infty}} \pi^{\infty}(x) \cdot P^{\infty}(x,y) \cdot C_{x,y}}{W/(M \cdot G^2)}$ and $\sum_{y \in S^{\infty}} \alpha(y) = 1$. Intuitively, $\alpha(y)$ is the stationary probability that the coalescence eventually occurs at state *y*. Eq. S27 implies that if ages and relative reproductive success in the initial sample are distributed according to $\alpha$, the expected *TMRCA*, in units of the generation time, is exactly $(M \cdot G)/W$. Note that we have made no asymptotic assumptions, i.e., this result is exact and holds even when the census size is very small. Also note that this result should hold in the more general context of the structured coalescent.

Next, we consider the expected *TMRCA* when the initial sample is not distributed according to $\alpha$ (e.g., when the alleles are sampled uniformly from a population at steady state). First, from Eq. S27 we see that

$$\left|e_x - \frac{M\cdot G^2}{W}\right| = \left|\sum_{y\in S^\infty} \alpha_y \cdot (e_x - e_y)\right| \le \sum_{y\in S^\infty} \alpha_y \cdot \left|e_x - e_y\right|. \quad \text{(S28)}$$

To bound the terms $\left|e_x - e_y\right|$, we consider the first visit of the chain to states $c$ or $y$. The time it takes to get from $x$ to $c$ can be partitioned to the time it takes to get to $c$ or $y$, plus the time to get from there to $c$, and thus

$$e_x = E\left(\tau^M_{\{y,c\}}\middle|X^M_0 = x\right) + P\left(X^M_{\tau^M_{\{y,c\}}} = y\middle|X^M_0 = x\right)e_y \le E\left(\tau^M_{\{y,c\}}\middle|X^M_0 = x\right) + e_y. \quad \text{(S29)}$$

To bound the term $E\left(\tau^M_{\{y,c\}}\middle|X^M_0 = x\right)$, we define a coupling of the chains $X^M_t$ and $X^\infty_t$, i.e. we define both chains on the same probability space, such that the chains have the exact same states until coalescence occurs. Formally, we assume that $X^M_0 = X^\infty_0$, and given $X^M_t, X^\infty_t$ such that $X^M_t \in \{X^\infty_t, c\}$, we define $X^M_{t+1}$ and $X^\infty_{t+1}$ as follows. First, $X^\infty_{t+1}$ is chosen with the appropriate probability conditioned on $X^\infty_t$. Second, if $X^M_t = c$ then $X^M_{t+1} = c$; if not, then $X^M_{t+1} = c$ with probability $C_{X^\infty_t, X^\infty_{t+1}}$, and $X^M_{t+1} = X^\infty_{t+1}$ otherwise. In this coupling, if $X^\infty_t = y$ then $X^M_t \in \{y, c\}$. It follows that

$$E\left(\tau^M_{\{y,c\}}\middle|X^M_0 = x\right) \le E\left(\tau^\infty_{\{y\}}\middle|X^\infty_0 = x\right). \quad \text{(S30)}$$

Defining $C = \max_{x,y\in S^\infty} E\left(\tau^\infty_{\{y\}}\middle|X^\infty_0 = x\right)$, we conclude that for any initial sample $x \in S^\infty$, the expected *TMRCA* in units of the generation time $G$ , $e_x/G$, satisfies

$$\left|\frac{e_x}{G} - \frac{M\cdot G}{W}\right| \le \frac{C}{G}. \quad \text{(S31)}$$

Since $C$ is defined on the chain $X^\infty_t$, it does not depend on the specific value of $M$. In other words, if we fix the age-structure (i.e., the breeding distribution $p$, the survival rates, and the distribution of relative reproductive success) and let the census size tend to infinity, the difference between the exact expected *TMRCA* and the asymptotic expectation, $M \cdot G/W$, remains bounded by $C/G$. Intuitively, the value of $C$ corresponds to the mixing time of the chain $X^\infty$, which is $O(1)$ and thus becomes negligible relative to the asymptotic expectation when $M$ is sufficiently large. In other words, Eq. S31 provides a bound on the difference between the asymptotic effective population size and its value for any given effective age class size $M$.

Previous works utilizing a similar framework relied on Möhle's lemma (5) to demonstrate weak convergence to Kingsman's coalescence process (3, 6). For example, considering a sample of size $n$ from the present population, and denoting the number of unique ancestors of the sample $t$ generations in the past by $Z_t$, the convergence to our Eq. 9 is stated as

$$Z_{\lfloor t/(\frac{G\cdot M}{W})\rfloor} \to R_t, \tag{S32}$$

where $(R_t)_{t\geq 0}$ is the standard Kingman coalescent for a sample of size $n$ (7), and the limit holds for any series of age-structured populations for which $G \cdot M/W \to \infty$. Although weak convergence does not generally imply convergence of moments, Möhle's lemma can be easily extended to show that convergence holds for the expected *TMRCA* of a sample of two alleles, i.e.,

$$\frac{E(TMRCA)}{G\cdot M/W} \to 1. \tag{S33}$$

While our result (Eq. S31) does not prove weak convergence (3) and is limited to a sample of two alleles, it provides proof for convergence of the first moment (i.e., it justifies Eq. 10 in the main text) and establishes tighter asymptotic rates of convergence compared to previous work (i.e., it proves that $\left|E(TMRCA) - \frac{G\cdot M}{W}\right| \leq C/G$). These results suggest that the asymptotic approximation is applicable even to very small populations.

#### 1.4 Reproductive variance

To recast our results for $N_e$ in terms of the total reproductive variance $V$, we first consider the case with non-overlapping generations in a haploid population of constant size, i.e., with Wright-Fisher sampling. We denote the number of offspring of the $i^{\text{th}}$ individual by $k_i$ and the census size by $N$. The expected number of offspring is 1, i.e., $\frac{1}{N}\sum_i k_i = 1$, and we denote the variance in number of offspring, which we also refer to as the reproductive variance, by $V = \frac{1}{N}\sum_i (k_i - 1)^2$. In the standard neutral model, without endogenous reproductive variance, $V = 1$. Since the probability that two distinct gametes descend from the same ancestor in the previous generation is $\sum_i \frac{k_i}{N} \cdot \frac{k_i - 1}{N-1} = \frac{V}{N-1}$, we find that the effective population size is

$$N_e = \frac{N-1}{V} \cong \frac{N}{V}, \tag{S34}$$

which is the expression derived by Wright (8) and presented in Eq. 12 of the main text.

To extend Eq. S34 to the case with overlapping generations, we consider the first two moments of an individual's number of offspring, $X$, throughout its lifetime. First, we note that an individual's number of offspring can be expressed as a sum over the number at each age, i.e., $X = \sum_a X_a$, where $X_a$ is the number of offspring at age $a$; $X_a = 0$ if the individual does not survive to that age. In these terms, the first two moments are

$$E(X) = \sum_a E(X_a) \text{ and } E(X^2) = \sum_a E(X_a^2) + 2\sum_{i<j} E(X_i \cdot X_j). \tag{S35}$$

Denoting the event of surviving to age ≥ $a$ by $S_a$, we note that

$$E\left(X_a^i\right) = Pr(S_a) \cdot E\left(X_a^i \middle| S_a\right) = \frac{M_a}{M_1} \cdot E\left(X_a^i \middle| S_a\right), \tag{S36}$$

The latter term, $E\left(X_a^i \middle| S_a\right)$, can be simplified further by conditioning on $\vec{r}$. Since the probability mass function of $\vec{r}$ conditional on $S_a$ is $f_a$,

$$E\left(X_a^i \middle| S_a\right) = E_{\vec{r}\sim f_a} E\left(X_a^i \middle| S_a, \vec{r}\right). \tag{S37}$$

Moreover, the distribution of $X_a$ conditional on $S_a$ and $\vec{r}$ is simply $(X_a|\vec{r}, S_a) \sim Bin(M_1, p_a \cdot r_a/M_a)$, and therefore

$$E(X_a|S_a) = E_{\vec{r}\sim f_a}\left(\frac{M_1 r_a p_a}{M_a}\right) = \frac{M_1 p_a}{M_a}$$

and

$$E(X_a^2|S_a) = E_{\vec{r}\sim f_a}\left(M_1\frac{r_a p_a}{M_a} + 2\binom{M_1}{2}\left(\frac{r_a p_a}{M_a}\right)^2\right) = \frac{M_1 p_a}{M_a} + 2\binom{M_1}{2}\left(\frac{p_a}{M_a}\right)^2 W_{a,a}. \tag{S38}$$

Substituting these expressions into Eq. S36, we find that

$$E(X_a) = p_a \text{ and } E(X_a^2) = p_a + \frac{M_1 - 1}{M_a} p_a^2 \cdot W_{a,a}. \tag{S39}$$

To calculate the remaining terms in Eq. S35, $E(X_i \cdot X_j)$ for $j > i$, we note that conditioning on $S_j$, and on $\vec{r}|S_j$,

$$E\left(X_i \cdot X_j\right) = Pr\left(S_j\right) \cdot E\left(X_i \cdot X_j \middle| S_j\right) = \frac{M_j}{M_1} \cdot E_{\vec{r}\sim f_j} E\left(X_i \cdot X_j \middle| S_j, \vec{r}\right). \tag{S40}$$

The latter term is easily calculated, since conditional on $S_j$ and $\vec{r}$, $X_i$ and $X_j$ are independent binomial variables, with $\left(X_i \middle| \vec{r}, S_j\right) \sim Bin(M_1, p_i \cdot r_i/M_i)$ and $\left(X_j \middle| \vec{r}, S_j\right) \sim Bin\left(M_1, p_j \cdot r_j/M_j\right)$, yielding

$$E\left(X_i \cdot X_j\right) = \frac{M_j}{M_1} \cdot E_{\vec{r}\sim f_j}\left(\frac{M_1^2 p_i p_j r_i r_j}{M_i M_j}\right) = \frac{M_1 p_i p_j W_{i,j}}{M_i}. \tag{S41}$$

Substituting the expressions from Eqs. S39 and S41 into Eq. S35 we find that

$$E(X) = 1 \text{ and } E(X^2) = 1 + M_1 \sum_i \frac{p_i^2 \cdot W_{i,i} + 2\sum_{j>i} p_i p_j W_{i,j}}{M_i} - \sum_i \frac{p_i^2 \cdot W_{i,i}}{M_i}. \tag{S42}$$

Assuming that the total population size is sufficiently large for the ratios $M_i/M_j$ and terms $W_{i,j}$ to be approximated as fixed, and for the higher order term $\sum_i \frac{p_i^2 \cdot W_{i,i}}{M_i}$ to be negligible, we find that

$$E(X) = 1 \text{ and } E(X^2) \cong 1 + \frac{M_1}{M} W, \tag{S43}$$

and therefore the total reproductive variance is

$$V = E(X^2) - E^2(X) = \frac{M_1}{M} W, \tag{S44}$$

which is Eq. 14 of the main text. These assumptions correspond to the standard practice of neglecting higher order terms in $1/N$ in models with non-overlapping generations. From Eqs. S18 and S44 we find that the effective population size is

$$N_e = (G \cdot M_1)/V, \tag{S45}$$

which is the same form as in the case without age-structure (Eq. S34), and the general form presented in Eq. 13 of the main text.

#### 1.5 Age-structure alone

Felsenstein used a different approach to solve the haploid model without endogenous reproductive variance, relying on the definition of the effective population size as the inbreeding effective number (2). To see that his results agree with ours (as well as with those of Sagitov and Jagers (3)), consider the case without endogenous reproductive variance, where Eq. S18 reduces to

$$N_e = MG = \frac{G}{\sum_i \frac{p_i^2 + 2\sum_{j>i} p_i p_j}{M_i}} = \frac{G}{\sum_i \frac{p_i}{M_i}(q_i + q_{i+1})}, \tag{S46}$$

where $q_i = \sum_{j \geq i} p_i$. Noting that $p_i(q_i + q_{i+1}) = (q_i - q_{i+1})(q_i + q_{i+1}) = q_i^2 - q_{i+1}^2$, we find that

$$N_e = \frac{G}{\sum \frac{p_i}{M_i}(q_i + q_{i+1})} = \frac{GM_1}{\sum \frac{M_1}{M_i}(q_i^2 - q_{i+1}^2)} = \frac{GM_1}{1 + \sum_i q_{i+1}^2 \left(\frac{M_1}{M_{i+1}} - \frac{M_1}{M_i}\right)}, \tag{S47}$$

where Felsenstein's functional form (p. 585 in (2)) is on the rightmost side.

To better understand the effect of age-structure on the effective population size, consider a simple example in which there is no endogenous reproductive variance, and no age-dependence in reproductive success. In other words, the only difference among individuals' numbers of offspring arise from the stochasticity of mortality and reproduction. In this case, the probability of having a parent of age $a$ is proportional to the size of the age class, i.e., $p_a = M_a/N$ where $N = \sum_a M_a$ is the census size. Following our derivations, the effective population size (Eq. 5 in the main text) then reduces to $N_e = \frac{G}{(2G-1)}N$, and if the generation time $G \gg 1$ then $N_e \approx \frac{1}{2}N$. In other words, the age structure reduces the effective population size to half of the census size.

#### 1.6 Upper bound on the effective population size

Here, we provide an upper bound for the effective population size of age-structured populations. With non-overlapping generations, the maximal effective population size equals the census population size, and it is attained when all individuals are equally likely to reproduce. In this case, endogenous reproductive variance reduces the effective population size below the census size (Eq. 12). In contrast, with age-structure, the maximal effective population size is attained when short-lived individuals are given a higher chance of reproducing while they live. In this case, we show that the effective population size can exceed the census size, but it is bound by the number of offspring per generation, $G \cdot M_1$. We also consider the conditions on $M_a$ and $p_a$ under which this upper bound can be attained, and describe the distributions of endogenous reproductive variances for which it is attained.

We begin by showing that the reproductive variance $V \geq 1$. Given that we have shown that in age-structured populations $N_e = G \cdot M_1/V$ (Eq. 13), showing that $V \geq 1$ establishes that $G \cdot M_1$ is an upper bound on $N_e$. To this end, we consider an individual's number of offspring, $X$, conditional on its relative reproductive success $\vec{r}$ and longevity $d$. Employing the notation of Section 1.4, $X = \sum_{i=1}^{d} X_i$, where the number of offspring at age $i$, $X_i \sim Bin(M_1, \frac{p_i r_i}{M_i})$, and the $X_i$s are independent of one another. It follows that

$$E(X|\vec{r}, d) = M_1 \sum_{i=1}^{d} \frac{p_i r_i}{M_i} \text{ and } Var(X|\vec{r}, d) = M_1 \sum_{i=1}^{d} \frac{p_i r_i}{M_i}\left(1 - \frac{p_i r_i}{M_i}\right) \cong E(X|\vec{r}, d), \quad \text{(S48)}$$

where the approximation for the variance becomes exact in the limit in which $M_i/M_j$ are held constant and the census population size goes to infinity. In other words, when the population size is sufficiently large, $X|(\vec{r}, d)$ is well approximated by a Poisson variable. From Eq. S49 and the law of total variance we have

$$V = E\,Var(X|\vec{r}, d) + Var\,E(X|\vec{r}, d) \cong E\,E(X|\vec{r}, d) + Var\,E(X|\vec{r}, d) = E(X) + Var\,E(X|\vec{r}, d)$$
$$= 1 + Var\,E(X|\vec{r}, d) \geq 1. \quad \text{(S48)}$$

Intuitively, Eq. S48 states that the in age-structured populations the distribution of the number of offspring is overdispersed due to stochasticity in longevity and endogenous reproductive variance. This implies that, in contrast to the case of non-overlapping generations, the number of offspring in age-structured populations is generally not well approximated by a Poisson variable. Eqs. 12 and S48 imply that

$$N_e \leq G \cdot M_1, \qquad \text{(S50)}$$

i.e., the effective population size is bound by the number of newborns per generation. This bound generalizes the bound $N_e \leq N$ in the case of non-overlapping generations.

Next, we consider an age-structured population with given values of $M_a$ and $p_a$, and ask which distributions of relative reproductive success, $f_a(\vec{r})$, maximize $N_e$, and what are the conditions on $M_a$ and $p_a$ for this maximum to equal the bound $G \cdot M_1$. Eq. S51 implies that maximizing $N_e$ is equivalent to minimizing $Var\ E(X|\vec{r}, d)$, and that the bound $G \cdot M_1$ is attained when $E(X|\vec{r}, d) = 1$ for any combination of $(\vec{r}, d)$ that occurs with non-zero probability. A distribution of $\vec{r}$ that minimizes $Var\ E(X|\vec{r}, d)$ can be explicitly constructed by the following algorithm:

1. Set $n = 1$. For each $d \geq 1$, initiate the $M_d - M_{d+1}$ vectors $\vec{r}$ of length $d$, corresponding to the $M_d - M_{d+1}$ individuals with longevity $d$, with zeros.
2. Choose the maximal $k \geq n$ at which the expression $\frac{M_1(q_n - q_{k+1})}{M_n - M_{k+1}}$ attains its minimum value $v$ (see Table S1 for the definition of $q$). For intuition, consider the case when $n = 1$: since there are $M_1 - M_{k+1}$ individuals with longevity $1 \leq d \leq k$, and $M_1(q_1 - q_{k+1})$ offspring that descend from parents in this range of ages, $v$ is an upper bound on the expected number of offspring that an individual with longevity in this range can have. We will assign vectors $\vec{r}$ such that $v$ is attained.
3. For $d = n, \dots, k$: Assign values $r_n, \dots, r_d$ to the individuals of longevity $d$, such that $\sum_{i=n}^{d} M_1 p_i \frac{r_i}{M_i} = v$, and under the constraint that $r_a$ over the $a^{\text{th}}$ age class should average to one. If $n > 1$, $r_1, \dots, r_{n-1}$ remain zero. Such an assignment always exists, but is not necessarily unique (Fig. S1).
4. If $k$ is smaller than the maximal longevity in the population, set $n = k + 1$ and return to step 2.

The bound $G \cdot M_1$ is attainable if and only if (iff) the algorithm requires exactly one step, which occurs iff

$$M_a \geq q_a \cdot M_1 \text{ for all } a \geq 1. \qquad \text{(S52)}$$

An example for an age-structured population that satisfies this condition, and for distributions of $\vec{r}$ given by the algorithm, are shown in Figs. S1A and S1B. The condition in Eq. S52 implies that

$$N_e \leq G \cdot M_1 = \sum_a q_a \cdot M_1 \leq \sum_a M_a = N \qquad \text{(S53)}$$

(note that $\sum_a q_a = G$ always holds), and thus that the effective population size cannot exceed the census size. In populations that do not satisfy the condition in Eq. S52, however, the effective population size is always smaller than $G \cdot M_1$ but can be larger than the census size (Fig. S1C).

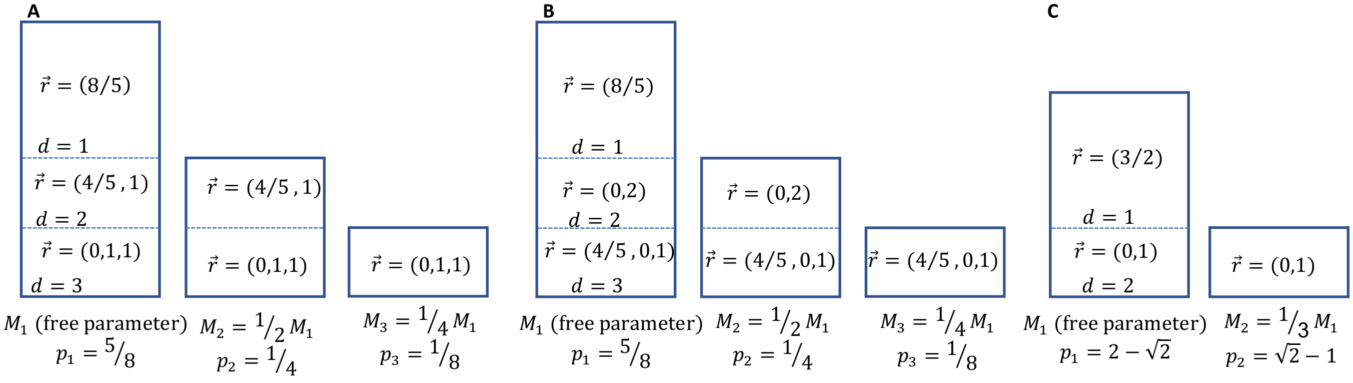

**Figure S1:** Maximal effective population sizes for specific age-structures. (A and B) An age structured population that satisfies the condition in Eq. S52 and thus has a maximal effective size equal to $G \cdot M_1$. In this example, $N_e = G \cdot M_1 = \frac{3}{2} \cdot M_1 < N = \frac{7}{4} \cdot M_1$. (A) and (B) show different distributions of $\vec{r}$ that derive from the algorithm and thus attain the maximal effective population size, illustrating that the construction is not unique. (C) An age-structure that does not satisfy the condition in Eq. S52. In this example the maximal effective size, $N_e = \frac{4+3\sqrt{2}}{6} \cdot M_1 \cong 1.37 \cdot M_1$, is strictly smaller than $G \cdot M_1 = \sqrt{2} M_1$, but larger than the census size, $\frac{4}{3} \cdot M_1$.

#### 1.7 Realized reproductive success

The model we considered so far allows for dependencies between endogenous but not realized reproductive success in different ages. Here we consider an alternative model that allows for such dependencies (also see (9)). Our model builds on the work of Sagitov and Jagers (3), who parametrize their model in terms of the distribution of the realized (rather than potential) number of offspring. We generalize their model to allow for any dependency between individuals' numbers of offspring in different ages. Importantly, we show that our formula for the effective population size (Eq. 13) holds under this model.

We assume that each newborn is assigned a vector $\vec{h} = (h_1, \dots h_{l(\vec{h})})$ of non-negative integers, such that it will survive to age $l(\vec{h})$, have $h_i$ offspring at age $i$, and have $S(\vec{h}) = \sum h_i$ offspring in total. The model's parameters consist of a set $H$ of $M_1$ such vectors, where $\sum_{\vec{h}\in H} S(\vec{h}) = M_1$ to ensure that the population size remains constant; this parametrization allows for general dependencies between individuals' numbers of offspring at different ages. At each time step, the $M_1$ vectors in $H$ are assigned to the $M_1$ newborns at random. The model is fully characterized by the set $H$. Notably, the age-class sizes are

$$M_a = \sum_{\vec{h}\in H:l(\vec{h})\geq a} 1 \tag{S54}$$

and the probability that a newborn descends from a parent of age $a$ is

$$p_a = \sum_{\vec{h}\in H:l(\vec{h})\geq a} h_a/M_1. \tag{S55}$$

The generation time is defined as $G = \sum i \cdot p_i$. We can also define an individual's relative reproductive success at a given age $a$ as the ratio of its realized success and the average realized success in that age-class, i.e.,

$$r_{\vec{h},a} = \frac{h_a}{\sum_{\vec{g}\in H:l(\vec{g})\geq a} g_a/M_a}; \tag{S56}$$

this definition is useful for comparing our main model with this one.

This model can be solved along the same lines we described in Sections 1.2 and 1.4. Here we provide only the main results, as the derivations are almost identical. First, the stationary distribution of $a$ (age) and $\vec{h}$ is

$$\epsilon(a,\vec{h}) = \frac{\sum_{\vec{h}\in H:l(\vec{h})\geq a}\sum_{i=a}^{l(\vec{h})} h_i}{M_1 \cdot G}. \tag{S57}$$

We rely on this distribution to solve for the stationary yearly rate of coalescence and corresponding effective population size. To this end, we define the effective age class size as $M \equiv (\sum w_a/M_a)^{-1}$, with weights

$$w_a = (\sum_{\vec{h}\in H:l(\vec{h})\geq a} p_a r_{\vec{h},a} (h_a - 1)/M_1 + 2\sum_{\vec{h}\in H:l(\vec{h})\geq a} p_a r_{\vec{h},a} (\sum_{i>a} h_i)/M_1)/W, \tag{S58}$$

where $W$ is defined such that these weights add up to 1. In these terms, the effective population size is well approximated by the same form as Eq. (10):

$$N_e \cong M \cdot G/W,$$

where the conditions for the approximation are the same as in Section 1.4. These results establish that $N_e$ takes the same form as it does for the model we described in the main text, although the definitions of $M$ and $W$ differ between the models.

We can also recast the results for this model in terms of total reproductive variance $V$. Calculating the total reproductive variance in this model is straightforward:

$$V = \frac{\sum_{\vec{h}\in H}\left(s(\vec{h})-1\right)^2}{M_1}. \quad \text{(S59)}$$

A simple rearrangement of terms in Eq. S57 then established that

$$N_e = G \cdot M_1/V,$$

which is Hill's formula and our Eq. 13. Thus, our main results apply under general dependencies between *realized* reproductive success in different ages.

Note, however, that in this model, the effective population size can exceed $G \cdot M_1$. The difference between models arises because in our main model, the distribution of the number of offspring is overdispersed compared to a Poisson distribution, whereas this model allows for the distribution to be under-dispersed. As an extreme example, consider a constant-sized population with non-overlapping generations, in which each individual has exactly one offspring; in this case, $V = 0$ and $N_e = \infty$ (coalescence never occurs).

### 2. Diploid Model

#### 2.1 Overview

Here we rigorously define and solve the diploid model with two sexes and endogenous reproductive variance, and derive formulas for the effective population sizes of X and autosomes. While the diploid model is more elaborate, the model and results follow along the same lines as we described for the haploid model. In Section 2.2 we detail the assumptions of the diploid model and introduce the notation required for the derivations that follow. In Section 2.3 we solve for the joint stationary distribution of the age and relative reproductive success of autosomal and X-linked alleles. We build on the joint stationary distribution to solving for the stationary per-generation coalescence rates and corresponding effective population sizes on X and autosomes. Since some of the explicit equations we derive are not presented in the main text, we briefly review them here.

Notably, to extend the haploid formula for the effective population size, $N_e = MG/W$ (Eq. 10), to the diploid case, we require explicit expressions for the effective age-class size $M$, generation time $G$, and $W$, corresponding to the X and autosomes. First, we define these measures for each sex in the same way that we did in the haploid model (i.e., as in Eqs. S16 and S17). We then define $G$ and $W$ for X and autosomes, as simple weighted averages over their values in males and females:

$$G_X = \frac{2}{3}G_F + \frac{1}{3}G_M \text{ and } G_A = \frac{1}{2}(G_M + G_F) \tag{S60}$$

and

$$W_X = \frac{2}{3}W_F + \frac{1}{3}W_M \text{ and } W_A = \frac{1}{2}(W_M + W_F) \tag{S61}$$

(Table 2 in the main text), where the weights reflect the relative number of generations that X and autosomal linked loci spend in males and females (see Table S2 for notation). The effective age class sizes on X and autosomes are defined as weighted harmonic averages. In the case without endogenous reproductive variance, they are defined as

$$\frac{1}{M_X} = \frac{1/3}{M_M} + \frac{2/3}{M_F} \text{ and } \frac{1}{M_A} = \frac{1/2}{M_M} + \frac{1/2}{M_F}. \tag{S62}$$

To account for sex-specific endogenous reproductive variances, the weights further account for the endogenous reproductive variances effect on the relative probability of coalescence in males and females,

$$\frac{1}{M_X} = \frac{1/3(W_M/W_X)}{M_M} + \frac{2/3(W_F/W_X)}{M_F} \text{ and } \frac{1}{M_A} = \frac{1/2(W_M/W_X)}{M_M} + \frac{1/2(W_F/W_X)}{M_F} \quad \text{(S63)}$$

(Table 2 in the main text). Using these definitions, the effective population size for the X and autosomes take the form

$$N_e^A = \frac{2G_A M_A}{W_A} \text{ and } N_e^A = \frac{2G_X M_X}{W_X}, \quad \text{(S64)}$$

where the factor 2, which is absent in the haploid case (Eq. 10), accounts for the effective number of age classes in the population (i.e., $G$ classes in the haploid population, but $2G$ classes in the case with two sexes). To translate these effective sizes into coalescence rates, we also account for ploidy, yielding per generation rates of $1/2N_e^A = W_A/4G_A M_A$ on autosomes and $1/(3/2)N_e^X = W_X/3G_X M_X$ on the X. Based on Eq. S65, the mutation rates on X and autosomes, and the standard forms for polymorphism levels, we obtain the following expression for the X/A polymorphism ratio:

$$\frac{E(\pi_X)}{E(\pi_A)} = \frac{3}{4} \cdot \frac{f(\mu_M/\mu_F)\cdot f(G_M/G_F)}{f\left(\frac{W_M/W_F}{M_M/M_F}\right)}. \quad \text{(S66)}$$

In Section 2.4, we recast the results for the effective population size (Eq. S67) and the X/A polymorphism ratio (Eq. S68) in terms of male and female reproductive variances. First, we show that the reproductive variances in males and females, $V_M$ and $V_F$, are given by

$$V_s = \frac{M_1}{\gamma_s}\frac{W_s}{M_s} - \frac{1-\gamma_s}{\gamma_s^2}, \quad \text{(S69)}$$

where the index $s$ corresponds to $M$ or $F$, and $\gamma_M$ and $\gamma_F$ are the proportions of males and females among newborns, respectively. Thus, this equation does not assume a sex ratio of 1. Rewriting Eq. S63 in terms of male and female reproductive variances we find that

$$N_e^X = \frac{4G_X M_1}{\frac{2}{3}\gamma_M V_M + \frac{4}{3}\gamma_F V_F + \frac{2\gamma_F}{3\gamma_M} + \frac{4}{3}\frac{\gamma_M}{\gamma_F}} \text{ and } N_e^A = \frac{4G_A M_1}{\gamma_M V_M + \gamma_F V_F + \frac{\gamma_F}{\gamma_M} + \frac{\gamma_M}{\gamma_F}}, \quad \text{(S70)}$$

where $M_1$ is the number of newborns of both sexes per year, and that

$$\frac{E(\pi_X)}{E(\pi_A)} = \frac{3}{4} \cdot \frac{f(\mu_M/\mu_F)\cdot f(G_M/G_F)}{f\left(\frac{\gamma_F/\gamma_M + \gamma_M V_M}{\gamma_M/\gamma_F + \gamma_F V_F}\right)}. \quad \text{(S71)}$$

These equations reduce to Eqs. 18 and 21 in the main text. In Section 2.5, we compare Eq. 18 (or Eq. S66) to Hill and Pollak's more complex expressions for age structure populations (see Introduction and (10-12)). We show that our formula is simpler in the general case in which the sex ratio at birth is not one. In Section 2.6, we recast our results in terms of reproductive success of alleles rather than individuals, in order to provide intuition for the differences in denominator between Hill's haploid formula, $N_e = G \cdot M_1/V$, and our extensions for diploids, e.g., $N_e = 4G_A M_1/(2 + V_A)$ for autosomes, assuming equal sex ratios at birth.

| Notation | Definition | Remarks |
|---|---|---|
| $p_{s,a}$ | Probability that a parent of sex $s$ is of age $a$ | $\Sigma_a p_{F,a} = \Sigma_a p_{M,a} = 1$ |
| $q_{s,a}$ | Probability that a parent of sex $s$ is of age $\geq a$ | $q_{s,a} = \Sigma_{i \geq a} p_{s,i}$ |
| $G_M, G_F$ | Male and female generation times | $G_s = \Sigma_a a \cdot p_{s,a}$ |
| $G_X, G_A$ | Generation times for X and autosomes | See Eq. S60 |
| $M_{s,a}$ | Number of individuals of sex $s$ and age $a$ | $M_{s,a+1} \leq M_{s,a}$ |
| $M_1$ | Number of newborns of both sexes per-year | |
| $\gamma_M, \gamma_F$ | Proportions of males and females among newborns | $\gamma_s = M_{s,1}/M_1$ |
| $\vec{r}$ | Relative reproductive success | |
| $f_{s,a}(\vec{r})$ | Proportion of individuals with relative reproductive success $\vec{r}$ among individuals of sex $s$ and age $a$ | |
| $g_{s,a}(\vec{r})$ | Given a newborn that descended from a parent of sex $s$ and age $a$, $g_{s,a}(\vec{r})$ is the probability that the parent has relative reproductive success $\vec{r}$ | $g_{s,a}(\vec{r}) = r_a \cdot f_{s,a}(\vec{r})$ |
| $\epsilon^X(s,a,\vec{r})$, $\epsilon^A(s,a,\vec{r})$. | Joint stationary probability of sex $s$, age $a$, and relative reproductive success $\vec{r}$ for the X and autosomes | See Eqs. S79 and S80 |
| $\epsilon^X_{s,a}, \epsilon^A_{s,a}$ | Marginal stationary distribution of sex $s$ and age $a$ for the X and autosomes | |
| $M_M, M_F$ | Effective male and female age-class sizes | See Eq. S91 |
| $M_X, M_A$ | Effective X and autosome linked age-class sizes | See Eq. S95 |
| $W_{s,i,j}$ | Expectation of $r_i \cdot r_j$ among individuals of sex $s$ conditional on surviving to age $a \geq j$ | Defined for $j \geq i$ |
| $W_M, W_F$ | Weighted averages of the $W_{M,i,j}$ and the $W_{F,i,j}$, respectively | See Eq. S90 |
| $W_X, W_A$ | Weighted averages of $W_M$ and $W_F$ for X and autosome linked loci | See Eq. S61 |
| $X_{s,a}, X_s$ | Random variables describing the number of offspring an individual of sex $s$ has at age $a$ or throughout life, respectively | |
| $V_M, V_F$ | Male and female reproductive variances (i.e., $V_s = V(X_s)$) | See Eq. S113 |
| $S_{s,a}$ | The event of a newborn of sex $s$ surviving to age $\geq a$ | |
| $f(x)$ | $f(x) \equiv (2x+4)/(3x+3)$ | |
| $\mu_M, \mu_F$ | Male and female expected mutation rates per generation | See Section 3 |
| $\mu_X, \mu_A$ | Expected mutation rates per generation on X and autosomes; $\mu_X = \frac{1}{3}\mu_M + \frac{2}{3}\mu_F$ and $\mu_A = \frac{1}{2}\mu_M + \frac{1}{2}\mu_F$ | |
| $X^X_m, X^A_m$ | The number of newborns carrying a random X or autosome linked allele $m$ | |
| $V^*_X, V^*_A$ | Reproductive variances of X and autosome linked alleles, respectively (i.e., $V^*_X \equiv V(X^X_m)$ and $V^*_A \equiv V(X^A_m)$) | See Eqs. S124 and S129 |

**Table S2:** Notation for the diploid model with two sexes, with parameters of the model in red.

#### 2.2 Assumptions and notation

We consider a panmictic, diploid population of constant size, with two sexes, and sex- and age-dependent mortalities, fecundities and reproductive variances. We measure age in years, and assume that the number of individuals of sex $s$ and age $a$, $M_{s,a}$, is constant. Specifically, the sizes of the newborn age classes, $M_{M,1}$ and $M_{F,1}$, may take any integer values, meaning that we do not assume that the sex-ratios at birth equals 1. More generally, the size of classes can vary between sexes, but for each sex they decrease with age, $\text{i.e.}, M_{s,a+1} \leq M_{s,a}$, reflecting sex- and age-specific mortalities. We further assume that age classes are partitioned according to individuals' age-dependent reproductive success. Namely, individuals are randomly assigned a vector $\vec{r}$ at birth, reflecting their expected relative reproductive success at each age (see below). We then assume that the number of individuals in the population of sex $s$, age $a$ and relative reproductive success $\vec{r}$, is constant and equal to $M_{s,a} \cdot f_{s,a}(\vec{r})$, where $f_{s,a}$ is the probability mass function of $\vec{r}$ among individuals of sex $s$ and age $a$. Individuals with the same value of $\vec{r}$ are chosen to survive to the next age class at random, i.e., there are no differences in viability, but $M_{s,a} \cdot f_{s,a}(\vec{r}) \geq M_{s,a+1} \cdot f_{s,a+1}(\vec{r})$ due to mortality, where rates of mortality can depend on the value of $\vec{r}$.

Sex and age dependent reproductive success is described backwards in time, in terms of the probability of an individual being chosen as a parent. Every newborn has a mother and a father, which are chosen independently. The probability that the parent of sex $s$ is of a given age is described by a discrete distribution $A_s = (p_{s,a})_{a=1}^{\infty}$, where the expectations $G_M = E(A_M)$ and $G_F = E(A_F)$ are the generation times for males and females, respectively. The average probability per individual of age $a$ is therefore $p_{s,a}/M_{s,a}$, which can be viewed as the fertility associated with that age and sex. The probability of being born to a specific parent of age $a$ and relative reproductive success $\vec{r}$ is $p_{s,a} \cdot \frac{r_a}{M_{s,a}}$, where $r_a$ is the $a$-th component of $\vec{r}$. The value of $r_a$ thus reflects an individual's expected (rather than realized) relative reproductive success.

Similar to the haploid case (cf. Section 1.1), our assumptions imply several requirements on the form of the probability mass functions $f_{s,a}$. First, requiring that the probability of a parent of sex $s$ being of age $a$ is $p_{s,a}$, implies that for any sex $s$ and age $a$, $E_{f_{s,a}}(r_a) = 1$. Second, requiring that

$M_{s,a} \cdot f_{s,a}(\vec{r}) \geq M_{s,a+1} \cdot f_{s,a+1}(\vec{r})$ implies that $f_{s,a}(\vec{r})/f_{s,a+1}(\vec{r}) \geq M_{s,a+1}/M_{s,a}$. Third, requiring that for any sex $s$ and age $a$, $M_{s,a} \cdot f_{s,a}(\vec{r})$ is an integer, implies that the probability mass functions $f_{s,a}$ are discrete and can only take values $i/M_{s,a}$ for $i = 0, 1, \dots, M_{s,a}$. While the latter requirement may appear to be highly restrictive, if we fix the ratios $M_{s,a}/M_{s',a'}$ and assume that the total population size is sufficiently large, we can relax this requirement and assume any continuous or discrete distributions $f_{s,a}$ that satisfy the first two requirements (by the same reasoning we applied to the haploid case in Section 1.2).

##### 2.3 Solution backwards in time

Here, we extend the derivations of Pollak (12) to account for endogenous reproductive variance. Tracing an allele backward in time, the sex $s_t$, age $a_t$ and relative reproductive success $\vec{r}_t$ of the individual $I_t$ carrying the allele $t$ years in the past defines a Markov chain, $(s_t, a_t, \vec{r}_t)$. To define the transition probabilities of the chain, we distinguish between two cases. First, if the allele is not carried by a newborn, i.e., if $a_t > 1$, then at time $t+1$ the individual carrying it was one year younger, and its sex $s$ and relative reproductive success $\vec{r}$ remain unchanged, i.e., $(s_{t+1}, a_{t+1}, \vec{r}_{t+1}) = (s_t, a_t - 1, \vec{r}_t)$ with probability 1. Second, if the allele is carried by a newborn, i.e., if $a_t = 1$, then the sex of the parent, $s_{t+1}$, is equally likely to be male or female if the allele is autosomal or if it is X-linked and the newborn was a female; if the allele is X-linked and the newborn was a male then the sex of the parent will be female with probability 1. Conditional on the parent's sex, $s_{t+1}$, its age $a_{t+1} = a$ with probability $p_{s_{t+1},a}$. The probability mass function of $\vec{r}_{t+1}$ conditional on $(s_{t+1}, a_{t+1})$, follows from Bayes' theorem, further conditioning on the fact that the parent, $I_{t+1} = I$, necessarily reproduced successfully

$$P(\vec{r}_I = \vec{r} | I_{t+1} = I, a_{t+1} = a)$$

$$= \frac{P(I_{t+1}=I|\vec{r}_{t+1}=\vec{r}, a_{t+1}=a) \cdot P(\vec{r}_I = \vec{r}|\ a_{t+1}=a)}{P(I_{t+1}=I)} = \frac{(r_a/M_{s,a}) \cdot f_{s,a}(\vec{r})}{\sum_{\vec{k}} (r_a/M_{s,a}) \cdot f_{s,a}(\vec{k})} = r_a \cdot f_{s,a}(\vec{r}). \quad \text{(S72)}$$

We denote this probability by $g_{s,a}(\vec{r}) \equiv r_a \cdot f_{s,a}(\vec{r})$, and conclude that when $a_t = 1$, $s_{t+1}$ is distributed as we described above and $P\big((a_{t+1}, \vec{r}_{t+1}) = (a, \vec{r}) \big| s_{t+1}\big) = p_{s,a} \cdot g_{s,a}(\vec{r})$.

$g_{s,a}$ is a proper probability mass function since $\sum_{\vec{r}} g_{s,a}(\vec{r}) = \sum_{\vec{r}} r_a \cdot f_{s,a}(\vec{r}) = 1$. Moreover, the parent's expected value of $r_a$ is $E_{\vec{r} \sim g_{s,a}}(r_a) = E_{\vec{r} \sim f_{s,a}}(r_a^2) = 1 + V_{\vec{r} \sim f_{s,a}}(r_a) \geq 1$. The latter inequality makes intuitive sense, as it implies that the allele is more likely to be descended from an individual that has higher than average relative reproductive success in its age class.

We rely on the transition probabilities to derive and solve recursions for the stationary probabilities, $\epsilon_A(s,a,\vec{r})$ and $\epsilon_X(s,a,\vec{r})$, of sex $s$, age $a$, and relative reproductive successes $\vec{r}$, of autosome and X linked alleles, respectively. For autosomal alleles

$$\epsilon^A(s,a,\vec{r}) = \epsilon^A(s,a+1,\vec{r}) + \left(\sum_{t,\vec{k}} \epsilon^A(t,1,\vec{k})\right) \cdot \frac{1}{2} p_{s,a} \cdot g_{s,a}(\vec{r}), \tag{S73}$$

where the first term corresponds to aging by one year and the second corresponds to parenting a newborn. For X linked alleles

$$\epsilon^X(s,a,\vec{r}) = \epsilon^X(s,a+1,\vec{r}) + \left(\frac{1}{2}\sum_{\vec{k}} \epsilon^X(F,1,\vec{k}) + \mathbb{I}_{s=F}\sum_{\vec{k}} \epsilon^X(M,1,\vec{k})\right) \cdot p_{s,a} \cdot g_{s,a}(\vec{r}), \tag{S74}$$

where $\mathbb{I}$ denotes an indicator function (i.e., $\mathbb{I}_{s=F}$ is 1 when $s = F$ and 0 otherwise), and, similar to the autosomal case, the first term corresponds to aging by one year and the second corresponds to parenting a newborn.

In order to solve these recursions, we first consider the marginal stationary distribution of age and sex, $\epsilon^A_{s,a} = \sum_{\vec{r}} \epsilon^A(s,a,\vec{r})$ for autosomes and $\epsilon^X_{s,a} = \sum_{\vec{r}} \epsilon^X(s,a,\vec{r})$. To this end, we sum the recursions over $\vec{r}$ to obtain recursions on the marginal distributions,

$$\epsilon^A_{s,a} = \epsilon^A_{s,a+1} + \left(\epsilon^A_{M,1} + \epsilon^A_{F,1}\right) \cdot \frac{1}{2} p_{s,a} \text{ and } \epsilon^X_{s,a} = \epsilon^X_{s,a+1} + \left(\frac{1}{2}\epsilon^X_{F,1} + \mathbb{I}_{s=F}\epsilon^X_{M,1}\right) \cdot p_{s,a}, \tag{S75}$$

where we also require that $\sum_{s,a} \epsilon^A_{s,a} = \sum_{s,a} \epsilon^X_{s,a} = 1$. These recursions were solved by Pollak (12) for the case without endogenous reproductive variance, yielding

$$\epsilon^A_{s,a} = q_{s,a}/2G_A,\ \epsilon^X_{M,a} = q_{M,a}/3G_X \text{ and } \epsilon^X_{F,a} = 2q_{F,a}/3G_X, \tag{S76}$$

where $q_{s,j} \equiv \sum_{j\geq a} p_{s,j}$ is the probability that a parent of sex $s$ is at least $j$ years old. Substituting these expressions into Eqs. S73 and S74, the recursions simplify to

$$\epsilon^A(s,a,\vec{r}) = \epsilon^A(s,a+1,\vec{r}) + \frac{1}{2G_a} p_{s,a} \cdot g_{s,a}(\vec{r}) \tag{S77}$$

for autosomes and

$$\epsilon^X(s,a,\vec{r}) = \epsilon^X(s,a+1,\vec{r}) + \frac{1+\mathbb{I}_{s=F}}{3G_X} \cdot p_{s,a} \cdot g_{s,a}(\vec{r}) \tag{S78}$$

for the X, where we further require that $\sum_{s,a,\vec{r}} \epsilon^A(s,a,\vec{r}) = \sum_{s,a,\vec{r}} \epsilon^X(s,a,\vec{r}) = 1$. The solution to these recursions is

$$\epsilon^A(s,a,\vec{r}) = \frac{1}{2G_A}\epsilon(s,a,\vec{r}) \tag{S79}$$

for autosomes and

$$\epsilon^X(s,a,\vec{r}) = \frac{1+\mathbb{I}_{s=F}}{3G_X}\epsilon(s,a,\vec{r}) \tag{S80}$$

for the X, where $\epsilon(s,a,\vec{r}) \equiv \sum_{j\geq a} p_{s,j} \cdot g_{s,j}(\vec{r})$.

The marginal stationary probability mass function of $\vec{r}$ follows,

$$\epsilon_{\vec{r}}^A = \sum_{s,a} \epsilon^A(s,a,\vec{r}) = \sum_{s,j} \frac{j \cdot p_{s,j}}{2G_A} \cdot g_{s,j}(\vec{r}) \tag{S81}$$

for autosomes, and

$$\epsilon_{\vec{r}}^X = \sum_{s,a} \epsilon^X(s,a,\vec{r}) = \sum_{s,j} \frac{(1+\mathbb{I}_{s=F}) \cdot j \cdot p_{s,j}}{3G_X} \cdot g_{s,j}(\vec{r}) \tag{S82}$$

for the X. These are proper probability mass functions since they are weighted averages of the probability mass functions $g_{s,j}$, since $\sum_{s,j} \frac{j \cdot p_{s,j}}{2G_A} = \sum_{s,j} \frac{(1+\mathbb{I}_{s=F}) \cdot j \cdot p_{s,j}}{3G_X} = 1$.

Similar to the haploid case, we rely on the stationary distribution to derive the probability of coalescence of two alleles. Consider the autosomal case first. For coalescence to occur at time *t* in the past, one of the alleles (A) would descend from the other (B) or both would descend from the same parental allele at that time (we provide examples for both scenarios in the haploid section). Specifically, if allele B is in an individual of sex *s*, age $a$ and relative reproductive success $\vec{r}$ at time $t$ (with probability $\epsilon^A(s,a,\vec{r})$), then allele A must be in a newborn at time $t-1$ (with probability $\epsilon_{M,1} + \epsilon_{F,1}$), having descended from the same individual carrying allele B (with probability $\frac{1}{2}p_{s,a} \cdot \frac{r_a}{M_a}$) and from allele B specifically (with probability ½). Summing over the individual's possible sexes, ages and reproductive success vectors, we obtain the probability

$$\sum_{s,a,\vec{r}} \epsilon^A(s,a,\vec{r}) \cdot \left(\epsilon_{M,1}^A + \epsilon_{F,1}^A\right) \cdot \frac{1}{2}p_{s,a} \cdot \frac{r_a}{2M_{s,a}} = \frac{1}{8(G_A)^2}\sum_{s,a} \frac{\sum_{j\geq a} p_{s,a} p_{s,j} \sum_{\vec{r}} r_a \cdot g_{s,j}(\vec{r})}{M_{s,a}}$$

$$= \frac{1}{8(G_A)^2}\sum_{s,a} \frac{\sum_{j\geq a} p_{s,a} p_{s,j} W_{s,a,j}}{M_{s,a}}, \tag{S83}$$

where, for $j \geq i$,

$$W_{s,i,j} \equiv E_{\vec{r}\sim f_{s,j}}\left(r_i \cdot r_j\right) = E_{\vec{r}\sim g_{s,j}}(r_i) = \sum_{\vec{r}} r_i \cdot g_{s,j}(\vec{r}) \tag{S84}$$

is the expectation of $(r_i \cdot r_j)$ over individuals of sex $s$ and age $j$. Further allowing for either allele or both to be the newborn, and using the inclusion-exclusion principal to subtract the probability

$$\left(\epsilon_{M,1}^A + \epsilon_{F,1}^A\right)^2 \sum_{s,a,\vec{r}} \left(\frac{1}{2} p_{s,a}\right)^2 \cdot \frac{r_a g_{s,a}(\vec{r})}{2M_{s,a}} = \frac{1}{8(G_A)^2} \sum_{s,a} \frac{p_{s,a} p_{s,a} W_{s,a,a}}{M_{s,a}} \quad \text{(S85)}$$

that both alleles were in a newborn prior to coalescence, the autosomal stationary coalescence rate per year is

$$\frac{1}{8(G_A)^2} \sum_{s,a} \frac{p_{s,a}^2 W_{s,a,a} + 2\sum_{j>a} p_{s,a} p_{s,j} W_{s,a,j}}{M_{s,a}}. \quad \text{(S86)}$$

The per generation coalescence rate (in terms of the autosomal generation time $G_A$) and corresponding effective population size are therefore

$$\frac{1}{2N_e^A} = \frac{1}{8 \cdot G_A} \sum_{s,a} \frac{p_{s,a}^2 W_{s,a,a} + 2\sum_{j>a} p_{s,a} p_{s,j} W_{s,a,j}}{M_{s,a}}. \quad \text{(S87)}$$

For the X, the stationary coalescence rate per year is

$$2\left(\epsilon_{M,1}^X + \frac{1}{2}\epsilon_{F,a}^X\right) \sum_{a,\vec{r}} \epsilon^X(F,a,\vec{r}) \cdot p_{F,a} \cdot \frac{r_a}{2M_{F,a}} + 2 \cdot \frac{1}{2}\epsilon_{F,a}^X \sum_{a,\vec{r}} \epsilon^X(M,a,\vec{r}) \cdot p_{M,a} \cdot \frac{r_a}{M_{M,a}}$$
$$-\left(\epsilon_{M,1}^X + \frac{1}{2}\epsilon_{F,1}^X\right)^2 \sum_{a,\vec{r}} p_{F,a}^2 \cdot \frac{r_a g_{F,a}(\vec{r})}{2M_{F,a}} - \left(\frac{1}{2}\epsilon_{F,1}^X\right)^2 \sum_{a,\vec{r}} p_{M,a}^2 \cdot \frac{r_a g_{M,a}(\vec{r})}{M_{M,a}}$$
$$= \frac{1}{9(G_X)^2} \sum_s (1 + \mathbb{I}_{s=F}) \sum_a \frac{p_{s,a}^2 W_{s,a,a} + 2\sum_{j>a} p_{s,a} p_{s,j} W_{s,a,j}}{M_{s,a}}, \quad \text{(S88)}$$

and the corresponding per generation coalescence rate, which defines the effective population size for the X, $N_e^X$, is

$$\frac{1}{(3/2)N_e^X} = \frac{1}{9G_X} \sum_s (1 + \mathbb{I}_{s=F}) \sum_a \frac{p_{s,a}^2 W_{s,a,a} + 2\sum_{j>a} p_{s,a} p_{s,j} W_{s,a,j}}{M_{s,a}} \quad \text{(S89)}$$

(defined in terms of the X-linked generation time $G_X$).

As outlined in Section 2.1, the effective population sizes, $N_e^X$ and $N_e^A$, can be rewritten in terms of the effective age class sizes, to obtain expressions that are analogous to Eq. 10 in the haploid case. To this end, the terms $G$, $W$ and $M$ in Eq. 10 need to be defined for the X and autosomes. First, we define these terms separately for males and females, by applying the haploid definitions. Specifically, we define

$$W_s = \sum_i p_{s,i}^2 W_{s,i,i} + 2\sum_{i<j} p_{s,i} p_{s,j} W_{s,i,j} \quad \text{(S90)}$$

as a weighted average of the $W_{s,i,j}$, and define

$$\frac{1}{M_s} = \sum_a \frac{w_{s,a}}{M_{s,a}} \quad \text{(S91)}$$

as a weighted harmonic average of the age classes sizes of sex *s*, with weights

$$w_{s,i} = (p_{s,i}^2 W_{s,i,i} + 2\sum_{j>i} p_{s,i} p_{s,j} W_{s,i,j})/W_s, \quad \text{(S92)}$$

where $\sum_a w_{s,a} = 1$. To extend the definitions of $G$, $W$ and $M$ to the X and autosomes, we define them as weighted averages over males and females. Specifically, $G$ and $W$ are defined as simple weighted averages,

$$G_A = \frac{1}{2}(G_M + G_F) \text{ and } G_X = \frac{2}{3}G_F + \frac{1}{3}G_M \quad \text{(S93)}$$

and

$$W_A = \frac{1}{2}(W_M + W_F) \text{ and } W_X = \frac{2}{3}W_F + \frac{1}{3}W_M. \quad \text{(S94)}$$

The effective age class size *M* for X and autosomes is defined as a weighted harmonic average,

$$\frac{1}{M_A} = \frac{1/2(W_M/W_A)}{M_M} + \frac{1/2(W_F/W_A)}{M_F} \text{ and } \frac{1}{M_X} = \frac{1/3(W_M/W_X)}{M_M} + \frac{2/3(W_F/W_X)}{M_F}. \quad \text{(S95)}$$

Expressing Eqs. S87 and S89 in these terms, we find that

$$N_e^A = \frac{2M_A G_A}{W_A} \text{ and } N_e^X = \frac{2M_X G_X}{W_X}, \quad \text{(S96)}$$

which is Eq. 15 in the main text. The factor 2, which is absent in the haploid case (Eq. S18), reflects the effective number of age classes (i.e., $G$ classes of size *M* in the haploid model, but *2G* classes in the diploid model with two sexes).

Assuming the standard expressions for neutral heterozygosity, $E(\pi_A) = 4N_e^A\mu_A$ and $E(\pi_X) = 3N_e^X\mu_X$ (see Section 3), and rearranging the expressions in Eq. S96, we find that

$$\frac{E(\pi_X)}{E(\pi_A)} = \frac{3}{4} \cdot \frac{f(\mu_M/\mu_F) \cdot f(G_M/G_F)}{f\left(\frac{W_M/W_F}{M_M/M_F}\right)}. \quad \text{(S97)}$$

When the mutation rate, age structure, and endogenous reproductive variance are identical in both sexes Eq. S97 reduces to the naïve neutral expectation of ¾. When these factors differ among sexes, Eq. S97 provides a simple expression for the effect of each factor.

#### 2.4 Reproductive variance

To recast our results for the effective population sizes in terms of total reproductive variances in males, $V_M$, and females, $V_F$, we follow the same steps as described for the haploid case (Section

1.4). First, we consider the case with non-overlapping generations in a diploid population of constant size, with $N_M$ males and $N_F$ females. We denote the total population size by $N \equiv N_M + N_F$, the proportions of males and females by $\gamma_s \equiv \frac{N_s}{N}$, and the number of offspring of the $i^{\text{th}}$ individual of sex $s$ by $k_i^s$. To maintain a constant population size, we require that the number of offspring arising from parents of each sex equals $N$, and therefore the sex-specific expectations are $E(k_i^s) = \frac{1}{N_s}\sum_i k_i^s = \frac{N}{N_s}$. We denote the sex-specific variances by $V_s \equiv V(k_i^s)$.

We are interested in the probability that two distinct alleles descend from the same allele in the previous generation, as this probability equals $1/2N_e^A$ for autosomes and $1/(3/2)N_e^X$ for the X. For autosomes, the probability that the two alleles descend from individuals of sex $s$ is ¼, the probability that they descend from the same individual of that sex is $\sum_{i=1}^{N_s} \frac{k_i^s}{N} \cdot \frac{k_i^s - 1}{N-1}$, and the probability that they descend from the same allele is 1/2, and therefore

$$\frac{1}{2N_e^A} = \frac{1}{8}\sum_s \sum_{i=1}^{N_s} \frac{k_i^s}{N} \cdot \frac{k_i^s - 1}{N-1}. \tag{S98}$$

Substituting $\sum_{i=1}^{N_s} \frac{k_i^s}{N} \cdot \frac{k_i^s - 1}{N-1} = \frac{\gamma_s}{N-1}\left(E\left({k_i^s}^2\right) - E(k_i^s)\right) = \frac{1}{N-1}(\gamma_s V_s + \frac{1}{\gamma_s} - 1)$ into Eq. S98, we find that

$$N_e^A = \frac{4(N-1)}{\gamma_M V_M + \gamma_F V_F + \gamma_F/\gamma_M + \gamma_M/\gamma_F} \cong \frac{4N}{\gamma_M V_M + \gamma_F V_F + \gamma_F/\gamma_M + \gamma_M/\gamma_F} \tag{S99}$$

(cf. (8)).

For the X chromosome, the probability that two alleles descend from individuals of sex $s$ depends on $\gamma_M$ and $\gamma_F$. However, as we go further backwards in time, this probability approaches 1/9 for both being male and 4/9 for both being female, regardless of $\gamma_M$ and $\gamma_F$. The probability that both alleles descend from the same individual of that sex is $\sum_{i=1}^{N_s} \frac{k_i^s}{N} \cdot \frac{k_i^s - 1}{N-1}$, and the probability that they descend from the same allele is ½ for females and 1 for males, and therefore

$$\frac{1}{(3/2)N_e^X} = \frac{1}{9}\sum_{i=1}^{N_M} \frac{k_i^M}{N} \cdot \frac{k_i^M - 1}{N-1} + \frac{1}{2} \cdot \frac{4}{9}\sum_{i=1}^{N_F} \frac{k_i^F}{N} \cdot \frac{k_i^F - 1}{N-1}, \tag{S100}$$

and thus

$$N_e^X = \frac{4(N-1)}{\frac{2}{3}\gamma_M V_M + \frac{4}{3}\gamma_F V_F + \frac{2}{3}\frac{\gamma_F}{\gamma_M} + \frac{4}{3}\frac{\gamma_M}{\gamma_F}} \cong \frac{4N}{\frac{2}{3}\gamma_M V_M + \frac{4}{3}\gamma_F V_F + \frac{2}{3}\frac{\gamma_F}{\gamma_M} + \frac{4}{3}\frac{\gamma_M}{\gamma_F}}. \tag{S101}$$

Assuming a sex ratio of 1 (i.e., $\gamma_M = \gamma_F = 1/2$), Eqs. S99 and S101 reduce to

$$N_e^A = \frac{4N}{2+\frac{1}{2}V_M+\frac{1}{2}V_F} \text{ and } N_e^X = \frac{4N}{2+\frac{1}{3}V_M+\frac{2}{3}V_F}. \tag{S102}$$

To extend these results to the case with overlapping generations, we consider the first two moments of an individual's number of offspring, $X_s$, throughout its lifetime. First, we note that an individual's number of offspring can be expressed as a sum over the number at each age, i.e., $X_s = \sum_a X_{s,a}$, where $X_{s,a}$ denotes the number of offspring at age $a$; and $X_{s,a} = 0$ if the individual does not survive to age $a$. In these terms, the first two moments are

$$E(X_s) = \sum_a E(X_{s,a}) \text{ and } E(X_s^2) = \sum_a E(X_{s,a}^2) + 2\sum_{j>i} E(X_{s,i} \cdot X_{s,j}). \tag{S103}$$

Denoting the event of surviving to age $\geq a$ by $S_{s,a}$, we note that

$$E\left(X_{s,a}^i\right) = Pr\left(S_{s,a}\right) \cdot E\left(X_{s,a}^i \middle| S_{s,a}\right) = \frac{M_{s,a}}{M_{s,1}} \cdot E\left(X_{s,a}^i \middle| S_{s,a}\right). \tag{S104}$$

The latter term, $E\left(X_{s,a}^i \middle| S_{s,a}\right)$, can be simplified further by conditioning on $\vec{r}$. Since the probability mass function of $\vec{r}$ conditional on $S_{s,a}$ is $f_{s,a}$,

$$E\left(X_{s,a}^i \middle| S_{s,a}\right) = E_{\vec{r}\sim f_{s,a}} E\left(X_{s,a}^i \middle| S_{s,a}, \vec{r}\right). \tag{S105}$$

Moreover, the distribution of $X_{s,a}$ conditional on $S_{s,a}$ and $\vec{r}$ is

$$\left(X_{s,a} \middle| \vec{r}, S_{s,a}\right) \sim Bin\left(M_1, p_{s,a} \cdot r_a / M_{s,a}\right), \tag{S106}$$

where $M_1 = M_{M,1} + M_{F,1}$ is the number of newborns of both sexes per-year, and therefore

$$E\left(X_{s,a} \middle| S_{s,a}\right) = E_{\vec{r}\sim f_{s,a}}\left(\frac{M_1 r_a p_{s,a}}{M_{s,a}}\right) = \frac{M_1 p_{s,a}}{M_{s,a}} \tag{S107}$$

and

$$E\left(X_{s,a}^2 \middle| S_{s,a}\right) = E_{\vec{r}\sim f_{s,a}}\left(M_1 \frac{r_a p_{s,a}}{M_{s,a}} + 2\binom{M_1}{2}\left(\frac{r_a p_{s,a}}{M_{s,a}}\right)^2\right) = \frac{M_1 p_{s,a}}{M_{s,a}} + 2\binom{M_1}{2}\left(\frac{p_{s,a}}{M_{s,a}}\right)^2 W_{s,a,a}.$$

Substituting these expressions into Eq. S104, we find that

$$E(X_{s,a}) = \frac{p_{s,a}}{\gamma_s} \text{ and } E\left(X_{s,a}^2\right) = \frac{p_{s,a}}{\gamma_s} + \frac{M_1 - 1}{M_{s,a}} \frac{p_{s,a}^2}{\gamma_s} \cdot W_{s,a,a}, \tag{S108}$$

where $\gamma_M$ and $\gamma_F$ are the proportions of males and females at birth (i.e., $\gamma_s = M_{s,1}/M_1$). To calculate the remaining terms in Eq. S103, $E(X_{s,i} \cdot X_{s,j})$ for $j > i$, we note that conditioning on $S_{s,j}$, and on $\vec{r}|S_{s,j}$,

$$E\left(X_{s,i} \cdot X_{s,j}\right) = P\left(S_{s,j}\right) \cdot E\left(X_{s,i} \cdot X_{s,j} \middle| S_{s,j}\right) = \frac{M_{s,j}}{M_{s,1}} \cdot E_{\vec{r}\sim f_{s,j}} E\left(X_{s,i} \cdot X_{s,j} \middle| S_{s,j}, \vec{r}\right). \tag{S109}$$

The latter term is easy to calculate: conditional on $S_{s,j}$ and $\vec{r}$, $X_{s,i}$ and $X_{s,j}$ are independent binomial variables: $(X_{s,i}|\vec{r},S_{s,j})\sim Bin(M_1, p_{s,i}\cdot r_i/M_{s,i})$ and $(X_{s,j}|\vec{r},S_{s,j})\sim Bin(M_1, p_{s,j}\cdot r_j/M_{s,j})$, and therefore

$$E(X_{s,i}\cdot X_{s,j}) = \frac{M_{s,j}}{M_{s,1}}\cdot E_{\vec{r}\sim f_{s,j}}\left(\frac{M_1^2 p_{s,i}p_{s,j}r_i r_j}{M_{s,i}M_{s,j}}\right) = \frac{M_1 p_{s,i}p_{s,j}W_{s,i,j}}{\gamma_s M_{s,i}}. \tag{S110}$$

Substituting the expressions from Eqs. S108 and S110 into Eq. S103 we obtain

$$E(X_s) = \frac{1}{\gamma_s} \text{ and } E(X_s^2) = \frac{1}{\gamma_s} + \frac{M_1}{\gamma_s}\sum_i \frac{p_{s,i}^2\cdot W_{s,i,i}+2\sum_{j>i}p_{s,i}p_{s,j}W_{s,i,j}}{M_{s,i}} - \sum_a \frac{p_{s,a}^2\cdot W_{s,a,a}}{\gamma_s M_{s,a}}. \tag{S111}$$

Assuming that the total population size is sufficiently large for the ratios $M_{s,i}/M_{t,j}$ and terms $W_{s,i,j}$ to be approximated as fixed, and for the higher order terms $\sum_a \frac{p_{s,a}^2\cdot W_{s,a,a}}{\gamma_s M_{s,a}}$ to be negligible, we find that

$$E(X_s) = \frac{1}{\gamma_s} \text{ and } E(X_s^2) \cong \frac{1}{\gamma_s} + \frac{M_1}{\gamma_s}\frac{W_s}{M_s}. \tag{S112}$$

The total reproductive variances of sex *s*, are therefore

$$V_s = E(X_s^2) - E^2(X_s) \cong \frac{M_1}{\gamma_s}\frac{W_s}{M_s} - \frac{1-\gamma_s}{\gamma_s^2}. \tag{S113}$$

From Eqs. S96 and S113, we obtain that

$$N_e^A = \frac{4G_A M_1}{\gamma_M V_M + \gamma_F V_F + \gamma_F/\gamma_M + \gamma_M/\gamma_F} \text{ and } N_e^X = \frac{4G_X M_1}{\frac{2}{3}\gamma_M V_M + \frac{4}{3}\gamma_F V_F + \frac{2}{3}\frac{\gamma_F}{\gamma_M} + \frac{4}{3}\frac{\gamma_M}{\gamma_F}}, \tag{S114}$$

where $G_A M_1$ and $G_X M_1$ are the total numbers of newborns per-generation, for autosomes and the X, respectively. Eq. S114, which is equivalent to Eq. 18 in the main text, generalizes Eqs. S99 and S101 to the case with age-structure.

Assuming that $E(\pi_A) = 4N_e^A\mu_A$ and $E(\pi_X) = 3N_e^X\mu_X$ (see Section 3), we find that

$$\frac{E(\pi_X)}{E(\pi_A)} = \frac{3}{4}\cdot\frac{f(\mu_M/\mu_F)\cdot f(G_M/G_F)}{f\left(\frac{\gamma_F/\gamma_M + \gamma_M V_M}{\gamma_M/\gamma_F + \gamma_F V_F}\right)}, \tag{S115}$$

which is Eq. 21 in the main text. When the sex ratio at birth is 1 (i.e. that $\gamma_M = \gamma_F = 1/2$), Eqs. S114 and S115 reduce to

$$N_e^A = \frac{4G_A M_1}{2+\frac{1}{2}V_M+\frac{1}{2}V_F},\ N_e^X = \frac{4G_X M_1}{2+\frac{1}{3}V_M+\frac{2}{3}V_F}, \text{ and } \frac{E(\pi_X)}{E(\pi_A)} = \frac{3}{4}\cdot\frac{f(\mu_M/\mu_F)\cdot f(G_M/G_F)}{f\left(\frac{2+V_M}{2+V_F}\right)}. \tag{S116}$$

##### 2.5 Hill and Pollak's results for age-structured populations

As we reviewed in the Introduction, Hill derived an expression for the effective population size of autosomes in age structured populations (Eq. 2) and Pollak derived a similar expression for the X (10-12). Here we show that these expressions apply to our extended model and compare them with our simpler expressions (Eqs. 18 and S114).

Hill and Pollak's results are cast in term of the variances and covariances of the number of male and female offspring of a parent of sex $s$, $X_{s,M}$ and $X_{s,F}$, respectively. Conditioned on $X_s$, the total number of offspring of either sex, $X_{s,M}$ and $X_{s,F}$ can be approximated by

$$X_{s,M}|X_s \sim Bin(X_s, \gamma_M) \text{ and } X_{s,F}|X_s, X_{s,M} = X_s - x_{s,M}, \tag{S117}$$

where this approximation neglects dependencies of the order of $1/M_1^2$ (i.e., the sexes of different offspring are not independent, as the total number of newborns of each sex is fixed). From this approximation, and the laws of total variance and total covariance, we attain

$$E(X_{s,t}) = \gamma_t/\gamma_s \text{ , } Var(X_{s,t}) = 1 - \gamma_s + \gamma_t^2 V_s \text{ and } Cov(X_{s,M}, X_{s,F}) = \gamma_M \gamma_F (V_s - 1/\gamma_s). \tag{S118}$$

Based on these relationships, we can substitute the sex ratio at birth and male and female reproductive variances in Eq. S114 by variances and covariances of $X_{s,t}$. Doing so for autosomes (as well as some arrangement of terms) yields

$$\frac{1}{N_e^A} = \frac{2 + Var(X_{M,M}) + \frac{M_{M,1}}{M_{F,1}} \cdot Cov(X_{M,M}, X_{M,F}) + \left(\frac{M_{M,1}}{M_{F,1}}\right)^2 \cdot Var(X_{M,F})}{16 M_{M,1} G_A} + \frac{2 + Var(X_{F,F}) + \frac{M_{F,1}}{M_{M,1}} \cdot Cov(X_{F,M}, X_{F,F}) + \left(\frac{M_{F,1}}{M_{M,1}}\right)^2 \cdot Var(X_{F,M})}{16 M_{F,1} G_A}, \tag{S119}$$

which is Hill's expression (1). A similar exercise for the X yields Pollak's expression (12) .

Our Eq. S114 simplifies these results considerably, as it circumvents the need to consider variances in the number of offspring of a specific sex separately, or the covariance between the number of sons and daughters. Notably, the covariance term, $Cov(X_{s,M}, X_{s,F})$, is quite complicated, because the numbers of sons and daughters both depend on the parent's survival and reproductive success.

#### 2.6 Allelic reproductive variance

Here we derive expressions for the effective population size of the X and autosomes in terms of reproductive success of alleles rather than of individuals. We show that substituting the allelic reproductive variance into the haploid expression for $N_e$ (Eqs. 13 and S45) yields the correct expression for autosomal alleles, whereas for the X this formulation applies only when the sex ratio at birth equals 1 (i.e., $\gamma_\mathrm{M} = \gamma_\mathrm{F} = 1/2$). These results provide intuition for the differences between our expression for $N_e$ in the haploid case (Eq. 13) and our expressions for X and autosomes (Eq. 18).

We begin with some motivation. We define the reproductive success of an allele as an individual's number of offspring that carry that allele, and denote the reproductive variance associated with X and autosome linked alleles by $V_X^*$ and $V_A^*$, respectively. In these terms, we might hope that our expression for $N_e$ in the haploid case (Eq. 13) would apply to the X and autosomes, i.e., that

$$\boldsymbol{\frac{3}{2}} \cdot N_e^X = G_X\left(\boldsymbol{\frac{3}{2}} \cdot M_1\right)/V_X^* \text{ and } \mathbf{2} \cdot N_e^A = G_A(\mathbf{2} \cdot M_1)/V_A^*, \tag{S120}$$

where the (bold) factors of 3/2 for the X and 2 for autosomes follow from considering the effective size for alleles rather than individuals on the left-hand side, and the number of newborn alleles rather than individuals on the right-hand side. Below, we show that assuming a sex ratio of 1 at birth, then

$$V_X^* = \frac{1}{4}\left(2 + \frac{1}{3}V_M + \frac{2}{3}V_F\right) \text{ and } V_A^* = \frac{1}{4}\left(2 + \frac{1}{2}V_M + \frac{1}{2}V_F\right), \tag{S121}$$

where the weights reflect the proportion of generations spent in males and females, and the additive factor 2 results from ploidy. Substituting these expressions into Eq. S120 we indeed obtain the correct effective population sizes, i.e.,

$$N_e^X = \frac{4G_X M_1}{2+\frac{1}{3}V_M+\frac{2}{3}V_F} \text{ and } N_e^A = \frac{4G_A M_1}{2+\frac{1}{2}V_M+\frac{1}{2}V_F}.$$

However, as we further show, this 'shortcut' yields the wrong answer for the X when the sex ratio at birth deviates from 1.

First, we calculate the allelic variances. Consider an allele $m$ carried by an individual $I_m$ of sex $s_m$. We define the allele's realized reproductive success as the number of $I_m$'s offspring who carry a copy of $m$, and denote it by $X_m^A$ when $m$ is autosomal and by $X_m^X$ when it is X-linked. We denote

$I_m$'s total number of offspring (whether they carry $m$ or not) by $X_I$. First consider an autosomal allele. Since each offspring of $I_m$ carries a copy of $m$ with probability ½, the conditional distribution $X_m^A|X_I \sim Bin(X_I, 1/2)$. From the law of total variance,

$$E(X_m^A) = \frac{1}{2}E(X_I) \text{ and } V(X_m^A) = \frac{1}{4}[E(X_I) + V(X_I)]. \quad \text{(S122)}$$

Further conditioning on the sex of the individual carrying the allele, $s_I$, we note that $E(X_I|s_I) = 1/\gamma_{s_I}$ (Eq. S112) and $V(X_I|s_I) = V_{s_I}$, where the individual $I_m$ is male with probability $\gamma_M$ and female with probability $\gamma_F$. Applying the law of total variance again, we find that

$$E(X_I) = 2 \text{ and } V(X_I) = \gamma_M V_M + \gamma_F V_F + \frac{(\gamma_M - \gamma_F)^2}{\gamma_M \gamma_F}. \quad \text{(S123)}$$

Substituting these expressions into Eq. S122, we find that

$$E(X_m^A) = 1 \text{ and } V_A^* \equiv V(X_m^A) = \frac{1}{4}[\gamma_M V_M + \gamma_F V_F + \frac{\gamma_M}{\gamma_F} + \frac{\gamma_F}{\gamma_M}]. \quad \text{(S124)}$$

When the sex ratio at birth is 1, and thus $\gamma_M = \gamma_F = 1/2$, Eq. S124) reduces to the autosomal part of Eq. S125. From Eqs. S114 and S124, we find that

$$N_e^A = \frac{G_A \cdot M_1}{V_A^*} \quad \text{(S126)}$$

for any sex-ratio. Given that the effective population sizes are defined by requiring coalescence rates of $1/N_e$ in haploids and $1/(2 \cdot N_e^A)$ in diploids, Eq. S126 is, in fact, analogous to Eq. S45.

Next, consider an X-linked allele. If the individual carrying the allele, $I_m$, is male, then only his female offspring will inherit the allele, and thus, $X_m^X|(s_I = M, X_I) \sim Bin(X_I, \gamma_F)$. Since $E(X_I|s_I = M) = 1/\gamma_M$ (Eq. S112) and $V(X_I|s_I) = V_{s_I}$, the law of total variance implies that

$$E(X_m^X|s_I = M) = \gamma_F/\gamma_M \text{ and } V(X_m^X|s_I = M) = \gamma_F^2 V_M + \gamma_F. \quad \text{(S127)}$$

The case in which $I_m$ is a female is similar to the autosomal case, and thus, $X_m^X|(s_I = F, X_I) \sim Bin(X_I, 1/2\,)$,

$$E(X_m^X|s_I = F) = \frac{1}{2\gamma_F} \text{ and } V(X_m^X|s_I = F) = \frac{1}{4}V_F + \frac{1}{4\gamma_F}. \quad \text{(S128)}$$

Given that there are $M_{M,1}$ X-linked alleles in newborn males and $2M_{F,1}$ in newborn females, the probability that an X-linked allele in a newborn is in a male is $\gamma_M/(1+\gamma_F)$ and the probability it is in a female is $2\gamma_F/(1+\gamma_F)$. Applying the law of total variance therefore implies that

$$E(X_m^X) = 1 \text{ and } V_X^* = Var(X_m^X) = \frac{\gamma_M \gamma_F^2}{1+\gamma_F} V_M + \frac{\gamma_F}{2(1+\gamma_F)} V_F + \frac{1+2\gamma_M\gamma_F}{2(1+\gamma_F)} + \frac{(1-2\gamma_F)^2}{2\gamma_M\gamma_F}. \quad \text{(S129)}$$

When the sex ratio at birth is 1, and thus $\gamma_M = \gamma_F = 1/2$, Eq. S129 reduces to the X related expression of Eq. S130. From Eqs. S114 and S129, we find that

$$N_e^X = \frac{G_X \cdot M_1}{V_X^*}, \tag{S131}$$

only holds when $\gamma_M = \gamma_F = 1/2$. Thus, the haploid result (Eqs. 13 and S45) applies to X-linked alleles only when the sex ratio at birth equals 1.

To explain why this result fails in the general case, consider the reproductive success of an X-linked allele in consecutive generations. As we have shown above, an allele's expected reproductive success is $\gamma_F/\gamma_M$ in males and $1/(2\gamma_F)$ in females (averaged over sexes the expectation is 1). Now consider the expected reproductive success in the next generation: if the allele was in a male in the previous generation it will necessarily be in a female, and the expected reproductive success of the offspring allele would be $1/(2\gamma_F)$; if the allele was in a female in the previous generation, the expected reproductive success is obtained by averaging over the sex of the offspring, and is $\frac{1}{2} + \gamma_F$. Thus, unless $\gamma_M = \gamma_F = 1/2$, the reproductive success of an X-linked allele will be negatively correlated between parents and offspring. This violates the assumption of the haploid model that the reproductive success of individuals and their offspring are independent.

### 3. Mutational process

Here we describe the assumptions on the mutational model and derive formulas for the expected levels of heterozygosity. To incorporate what has recently been revealed about the dependencies of mutation rates on sex and age (e.g., (13-16)), we allow for mutation rate in the diploid model to depend on sex and age. Namely, we assume that the number of de novo mutations that a parent of sex $s$ and age $a$ bequeaths to its newborn is a random variable with expectation $\mu_{s,a}$ per base pair. Since mutation rates can vary with sex and age, the mutation rates per generation in males and females depend on the distributions of their breeding ages (i.e. $A_M$ and $A_F$, which were defined in Section 2). We denote the expected mutation rate per generation in males by $\mu_M = E_{A_M}(\mu_{M,a}) = \sum_a p_{M,a} \cdot \mu_{M,a}$ and the expected rate in females by $\mu_F = E_{A_F}(\mu_{F,a})$. The average rates on the autosomes and the X are given by $\mu_A = \frac{1}{2}(\mu_M + \mu_F)$ and $\mu_X = \frac{2}{3}\mu_F + \frac{1}{3}\mu_M$ (Table 2). For the haploid model, we assume the expected number of mutations $\mu_a$ to be dependent of age and define the per generation rate as $\mu = E_A(\mu_a)$. In the special case in which the parameters $\mu_{s,a}$ (or the $\mu_a$ in the haploid case) depend linearly on age, these expectations will depend only on the expected generation times $G_M$ and $G_F$, i.e., they are insensitive to higher moments of the distributions of breeding ages in males and females. As we show below, higher moments of the distributions of mutation rates per generation do not affect our results, which is how we avoid any further assumptions about these distributions.

The standard expressions for heterozygosity (e.g., $E(\pi_A) = 4N_e^A\mu_A$) are usually derived assuming that the genealogical and mutational processes are independent (17). This assumption is violated in our case, because both the time to the most recent common ancestor and the number of accumulated mutations depend on the ages of the individuals along the lineage. To derive the expected autosomal heterozygosity $E(\pi_A)$ under these conditions, we track alleles $A$ and $B$ backwards in time. Let $X_i$ denote the number of mutations occurring on the lineage leading from allele $A$ in the $i^{th}$ generation and $T$ denote the number of generations until the alleles coalesce. The number of mutations on the lineage leading to allele $A$ is then $\sum_{i=1}^{T} X_i$. Although $X_i$ and $T$ are dependent variables, Wald's equation (18) implies that $E(\sum_{i=1}^{T} X_i) = E(T) \cdot E(X_i)$ (to see that Wald's equation holds, note that the indicator function $\mathbb{I}_{T\geq n}$ is independent of $X_n$, since the first depends on the sexes and ages in the first $n - 1$ generations, and the second on the $n^{\text{th}}$ generation). We have already shown that $E(T) = 2N_e^A$. Since $E(X_i|s_i, a_i) = \mu_{s,a}$ (where $s_i$ and $a_i$ are the sex and age in the $i^{\text{th}}$ generation), it follows that $E(X_i) = E(\mu_{s,a}) = \mu_A$. We conclude that the lineage leading to allele A has on average $E(\sum_{i=1}^{T} X_i) = 2N_e^A \cdot \mu_A$ mutations and therefore $E(\pi_A) = 4N_e^A \mu_A$. A similar argument shows that for the haploid model $E(\pi) = 2N_e \mu$.

This argument requires modification for the X-chromosome, because the sexes $s_i$ and $s_{i+1}$ in consecutive generations along the lineage are dependent variables, leading to a dependence between $X_{i+1}$ and $s_i$, in violation of the conditions for Wald's equation. Instead, we define $T$ as the number of females on the lineage until the coalescence occurs, and define $X_i$ as the number of mutations between the $i^{\text{th}}$ and $i + 1$ females on the lineage. Under this definition, Wald's equation holds and $E(\pi_X) = 2E(\sum_{i=1}^{T} X_i) = 2E(X_i)E(T)$. It is then easy to show that $E(X_i) = \frac{3}{2}\mu_X$ and $E(T) = N_e^X$, implying that $E(\pi_X) = 3N_e^X \mu_X$.

### 4. Life history and population size that change over time

Thus far we considered models with constant population size and life history parameters. Here we extend our results to models in which population size and life history traits change over time. Specifically, we consider models with piecewise-constant age-structures, endogenous reproductive variances and population sizes. We rely on our results showing that with constant population sizes, the coalescence process with age structure is well approximated by Kingsman's coalescence process with non-overlapping generations, with the appropriate parameters (i.e., effective population sizes and time units). This allows us to approximate the coalescence rates per year in each time interval with constant parameters. We then derive simple recursions for expected heterozygosities on the X and autosomes at any point in time (Section 4.2), rely on these recursions to solve the example discussed in the main text (Eq. 24 and Fig 2), and specify how existing coalescence simulators can be used in order to account for life history effects (Section 4.3). The approximations we detail can be also be applied to models with multiple populations and piecewise-constant migration rates.

#### 4.1 Model

We measure time in years, backwards from the present, $t = 0$, and assume we are given $n$ time intervals, where the $i$-th interval is $[T_{i-1}, T_i)$, and $0 = T_0 < T_1 < \cdots < T_n = \infty$. The effective population sizes, generation times, and mutation rates, for the X and autosomes are defined as before, and are assumed to be constant in each time interval, where we denote their values in the $i$-th interval with an addition index $i$.

It may sometimes be useful to specify the model in terms of different, equivalent sets of parameters. For example, in Amster et al. (19), we rely on estimates of the autosomal effective population size, $N_e^{A,i}$, for a given set of time intervals, $i = 1, \dots, n$, which were inferred for human populations, assuming a constant autosomal generation time of $G_A = 30$ years. We then express the effective population sizes and generation times on the X in these intervals in terms of the sex ratios of generation times, $(G_M/G_F)_i$, and reproductive variances, $[(\gamma_M V_M + \gamma_F/\gamma_M)/(\gamma_F V_F + \gamma_M/\gamma_F)]_i$, for $i = 1, \dots, n$, as

$$N_e^{X,i} = \frac{f((G_M/G_F)_i)}{f([(\gamma_M V_M + \gamma_F/\gamma_M)/(\gamma_F V_F + \gamma_M/\gamma_F)]_i)} \cdot N_e^{A,i}, \quad \text{(S132)}$$

(relying on Eq. 20) and

$$G_X^i = 2G_A[1/3 + 1/(1 - (G_M/G_F)_i)]. \quad \text{(S133)}$$

We also rely on a model that describes human maternal and paternal mutation rates as a function of their respective generation times (20), and therefore specify the mutation rates per generation on the X and autosomes in time interval *i* as

$$\mu_A^i = \mu_A(G_A, (G_M/G_F)_i) \text{ and } \mu_X^i = \mu_X(G_A, (G_M/G_F)_i). \quad \text{(S134)}$$

#### 4.2 A recursion for heterozygosity on X and autosomes

Provided the effective population size, generation times, and mutation rates in each time interval for the X and autosomes, we can write down a simple recursion for the expected heterozygosities at present, $\pi_X$ and $\pi_A$. We first consider the autosomal case, and denote the expected heterozygosity at time $t$ by $\pi_A(t)$. The *n*-th time interval, $[T_{n-1}, \infty)$, is infinitely long with constant effective population size and mutation rate, and therefore

$$\pi_A(T_{n-1}) = 4N_e^{A,n}\mu_A^n. \quad \text{(S135)}$$

Next, we assume that we know that we know $\pi_A(T_i)$ and solve for $\pi_A(T_{i-i})$. To this end, we denote the event of two alleles sampled at time $T_{i-i}$ coalescing in the *i*-th time interval (i.e., until time $T_i$) by $E$, and its complement, i.e., that they do not coalesce, by $E^C$. Under the infinite sites assumption, the heterozygosity at time $T_{i-i}$ is then

$$\pi_A(T_{i-i}) = \mathrm{P}(E) \cdot \pi_A(T_{i-i}|E) + \mathrm{P}(E^C) \cdot \pi_A(T_{i-i}|E^C)$$

$$= \mathrm{P}(E) \cdot \left(2 \cdot \mathrm{E}(t_{MRCA}|E) \cdot \mu_A^i/G_A^i\right) + \left(1 - \mathrm{P}(E^C)\right)\left(\pi_A(T_i) + 2(T_i - T_{i-i}) \cdot \mu_A^i/G_A^i\right), \quad \text{(S136)}$$

where P denotes probability, and $t_{MRCA}$ denotes the time to the most recent common ancestor of the aforementioned sample. Approximating the time to coalescence in the *i*-th interval with an exponential distribution with rate $1/2G_A^i N_e^{A,i}$ (where $G_A^i$ is included for the process to be describe in years rather than generations), we find that

$$\mathrm{P}(E) = 1 - \exp\left(-\frac{T_i - T_{i-1}}{2G_A^i N_e^{A,i}}\right), \quad \text{(S137)}$$

and

$$E(t_{MRCA}|E) = 2G_A^i N_e^{A,i} - (T_i - T_{i-1})\frac{1 - \mathrm{P}(E)}{\mathrm{P}(E)}. \quad \text{(S138)}$$

Substituting these expressions into Eq. S136, we find that

$$\pi_A(T_{i-1}) = \left(1 - \exp\left(-\frac{T_i - T_{i-i}}{2G_A^i N_e^{A,i}}\right)\right) \cdot 4N_e^{A,i}\mu_A^i + \exp\left(-\frac{T_i - T_{i-i}}{2G_A^i N_e^{A,i}}\right) \cdot \pi_A(T_{i+1}). \tag{S139}$$

By the same token, we find that for the X:

$$\pi_X(T_{n-1}) = 3N_e^{X,n}\mu_X^n, \tag{S140}$$

and

$$\pi_X(T_{i-1}) = \left(1 - \exp\left(-\frac{T_i - T_{i-i}}{(3/2)G_X^i N_e^{X,i}}\right)\right) \cdot 3N_e^{X,i}\mu_X^i + \exp\left(-\frac{T_i - T_{i-i}}{(3/2)G_X^i N_e^{X,i}}\right) \cdot \pi_X(T_{i+1}). \tag{S141}$$

Eqs. S135 and S139-S141 can be solved recursively for $\pi_A$ and $\pi_X$ (i.e., for $\pi_A(T_0)$ and $\pi_X(T_0)$), respectively. The same recursions can be used to solve for $\pi_A(t)$ and $\pi_X(t)$ at any time $t$, where for $t$ in the $i$-th interval, we solve the same recursion until time $T_i$, and for the last step, we replace $T_{i-1}$ by $t$ in Eqs. S139 and S141.

##### 4.3 Simulations

The coalescence process on the X and autosomes accounting for life history effects can be simulated using standard tools. For example, to use *ms* (21), one can use per-generation coalescence rates (e.g., $1/2N_e^{A,i}$ and $1/(3/2)\,N_e^{X,i}$ for intervals $i = 1, \dots, n$, in the aforementioned model), and convert the duration of time intervals into units of generations, with the appropriate generation time (e.g., $(T_i - T_{i-i})/G_A^i$ and $(T_i - T_{i-i})/G_X^i$ for intervals $i = 1, \dots, n$, in the aforementioned model).

Simulating the mutational process requires a custom tool, as the standard ones assume fixed mutation rates, whereas we assume rates can change. It is, however, straightforward to implement such a tool, e.g., taking trees generated by ms (i.e., with the -T flag) as input, and incorporating piecewise constant mutation rates. We provide such a tool in https://github.com/sellalab/XA_poly.
